# Global genomics of the man-o’-war (*Physalia*) reveals biodiversity at the ocean surface

**DOI:** 10.1101/2024.07.10.602499

**Authors:** Samuel H. Church, River B. Abedon, Namrata Ahuja, Colin J. Anthony, Diego A. Ramirez, Lourdes M. Rojas, Maria E. Albinsson, Itziar Álvarez Trasobares, Reza E. Bergemann, Ozren Bogdanovic, David R. Burdick, Tauana J. Cunha, Alejandro Damian-Serrano, Guillermo D’Elía, Kirstin B. Dion, Thomas K. Doyle, João M. Gonçalves, Alvaro Gonzalez Rajal, Steven H. D. Haddock, Rebecca R. Helm, Diane Le Gouvello, Zachary R. Lewis, Bruno I. M. M. Magalhães, Maciej K. Mańko, Alex de Mendoza, Carlos J. Moura, Ronel Nel, Jessica N. Perelman, Laura Prieto, Catriona Munro, Kohei Oguchi, Kylie A. Pitt, Amandine Schaeffer, Andrea L. Schmidt, Javier Sellanes, Nerida G. Wilson, Gaku Yamamoto, Eric A. Lazo-Wasem, Chris Simon, Mary Beth Decker, Jenn M. Coughlan, Casey W. Dunn

## Abstract

The open ocean is a vast, highly connected environment, and the organisms found there have been hypothesized to represent massive, well-mixed populations. Of these, the Portuguese man-o’-war (*Physalia*) is uniquely suited to dispersal, sailing the ocean surface with a muscular crest. We tested the hypothesis of a single, panmictic *Physalia* population by sequencing 133 genomes, and found five distinct lineages, with multiple lines of evidence showing strong reproductive isolation despite range overlap. We then scored thousands of citizen-science photos and identified four recognizable morphologies linked to these lineages. Within lineages, we detected regionally endemic subpopulations, connected by winds and currents, and identified individual long-distance dispersal events. We find that, even in these sailing species, genetic variation is highly partitioned geographically across the open ocean.

**Summary:** The open ocean is a vast and highly connected environment. The organisms that live there have a significant capacity for dispersal and few geographic boundaries to separate populations. Of these, the Portuguese man-o’-war or bluebottle (genus *Physalia*) is uniquely suited to long-distance travel, using its gas-filled float and muscular crest to catch the wind and sail the sea surface. *Physalia* are distributed across the globe, and like many pelagic organisms, have been hypothesized to represent a massive, well-mixed population that extends across ocean basins. We tested this hypothesis by sequencing whole genomes of 133 samples collected from waters of over a dozen countries around the globe. Our results revealed five distinct lineages, with multiple lines of evidence indicating strong reproductive isolation, despite regions of range overlap. We combined these data with an independent dataset of thousands of images of *Physalia* uploaded to the citizen-science website inaturalist.org, which we scored for morphological characters including sail size, tentacle arrangement, and color. From these images, we identified four recognizable morphologies, described their geographical distribution, and linked them to four of the lineages identified with genomic data. We conclude there are at least four species, three of which correspond to species proposed by scientists in the 18th and 19th centuries: *P. physalis*, *P utriculus*, and *P. megalista*, along with one as yet unnamed species *Physalia* sp. from the Tasman Sea. Within each species, we observe significant population structure, with evidence of persistent subpopulations at a regional scale, as well as evidence for individual long-distance dispersal events. Our findings indicate that, instead of one well-mixed, cosmopolitan species, there are in fact multiple *Physalia* species with distinct but overlapping ranges, each made up of regionally endemic subpopulations that are connected by major ocean currents and wind patterns.

## Introduction

The open ocean has few geographic barriers that might limit connectivity (*1*). The organisms that live there often have strong dispersal potential (*2*) and massive effective population sizes (*3*), contributing to the assumption that populations are predominantly well-mixed, even at a global scale. However, a series of recent studies have found evidence for population structure in the open ocean, despite the absence of geographic barriers (*4*–*6*). Those studies challenge expectations of uninterrupted gene flow and bolster claims that open-ocean diversity has routinely been underestimated (*7*).

Studies of oceanic population structure have largely focused on benthic and planktonic species (either holoplanktonic or planktonic in the larval stage), meaning far less is known about populations that live at or near the ocean surface (*8*), collectively termed neuston (*9*). The surface ecosystem represents a unique biological environment, and the physical processes at play at the air-water interface (e.g. winds, surface currents) have distinct potential to mediate dispersal (*10*). At the same time, the ocean surface ecosystem is imperiled by plastics and pollutants that aggregate there, as well as by efforts to clean pollutants at a large scale (*11*). A common but unproven justification for potentially destructive clean-up efforts is that there is relatively little diversity at the ocean surface, and the organisms present there have robust population sizes (*12*). It is urgent that we evaluate this claim by examining genetic diversity at the surface to build informed strategies moving forward (*13*).

Bluebottles or Portuguese man-o’-war, cnidarians in the genus *Physalia*, present a compelling test case for exploring open-ocean population structure. They are among the few invertebrates to utilize wind-powered movement, sailing the ocean surface with a muscular crest, and they are the largest to do so, making them particularly capable of long-distance dispersal. There is only one species of *Physalia* currently recognized, with a hypothesized population that extends across the Atlantic, Indian, Pacific, and Southern Oceans (*14*–*16*). However, a recent study, analyzing marker genes from samples around New Zealand, found preliminary evidence of substantial genetic variation, even within a relatively small geographic area (*17*). An analysis of genomic variation in *Physalia* therefore represents a prime test for the existence of a globally panmictic population, targeting a widespread taxon with a significant capacity for long-distance dispersal (*18*).

*Physalia* populations are potentially influenced by the dynamics both at and below the ocean surface. Reproduction occurs below the surface, as reproductive structures (gonodendra) separate from the main body, sink, and release gametes into the water column (*19*). Following fertilization, juvenile *Physalia* return to the surface using specialized gas-producing tissues to inflate their nascent float (*20*). Growth occurs through the addition of asexually-budded, clonal bodies (called zooids) that remain integrated to one another through shared nervous and gastric systems, similar to a colony of coral, but in *Physalia* these bodies perform specialized functions (e.g., reproduction, prey capture, digestion) (*14*). Mature *Physalia* colonies are key predators within the neuston assemblage (*21*), extending their tentacles up to tens of meters into the water column to kill and retrieve fish (*22*). They additionally serve as prey within the neuston (*9*, *23*) and shoreline ecosystems (*24*), since onshore winds can blow these colonies onto beaches, often in large numbers (*25*, *26*). Given their potent sting, near-shore arrivals present a medical risk to humans and affect tourism via beach closures. These impacts create an additional need to understand the factors influencing their dispersal and distribution (*27* –*29*).

Variation in colony size is associated with ocean region (e.g., Atlantic specimens are typically the largest). Two alternative hypotheses can explain this pattern (*14*, *16*): [1] the large *Physalia* in certain parts of the world represent the oldest colonies – ones that sailed in from elsewhere, or [2] there are distinct populations in different regions that reach different sizes at maturity. New technologies make it possible to distinguish between these hypotheses, including increased efficiency of next-generation sequencing that has made it feasible to collect genomic data despite their large genome size (estimated at 2-3Gb (*30*)). In addition, participatory science on the internet has generated thousands of images of *Physalia* from beaches and waters around the world (Fig. 1B). In this study, we evaluate the population structure and diversity of *Physalia* by evaluating two independent datasets of *Physalia* diversity: [1] whole genome sequencing of 133 specimens, and [2] morphological data from more than 4,000 images submitted via participatory science to the natural history website inaturalist.org. We test for evidence of multiple species associated with distinct morphologies, describe their ranges and distributions, and analyze the spatiotemporal dynamics within each lineage.

**Figure 1:**
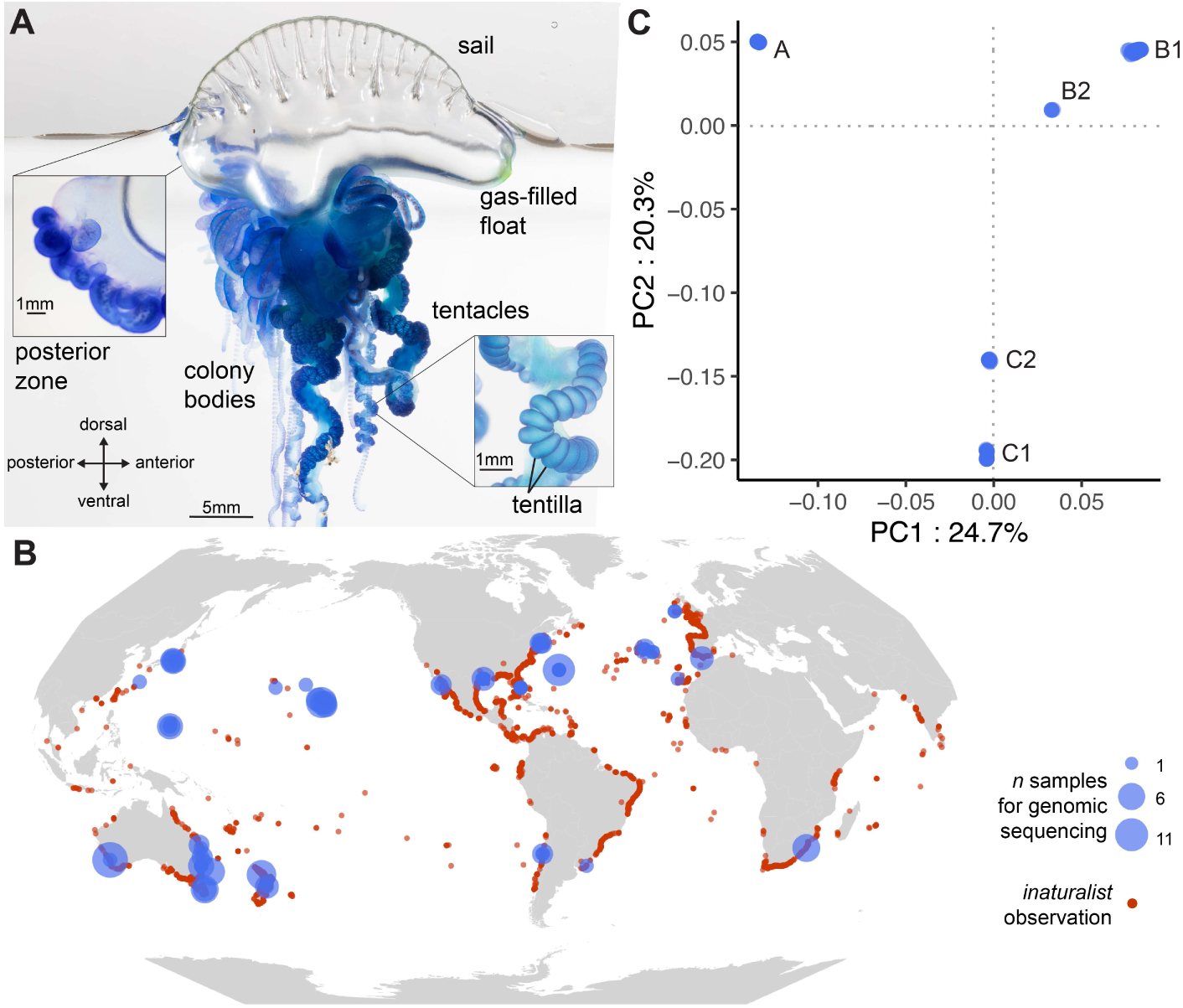
Anatomy, distribution, and genomic variation of *Physalia*. A, *Physalia* colonies comprise a muscular sail attached to a gas-filled float which maintains the mature animal at the surface of the water. Colony bodies (zooids), including those specialized for feeding (gastrozooids), prey capture (palpons with tentacles), and reproduction (gonozooids) are added to the float via asexual reproduction at growth zones. Tentacles drape below the float to trap, sting, and retrieve fish using batteries of stinging capsules contained in tentilla. Photos of *Physalia* sp. C2, specimens YPM IZ 111236 (main), YPM IZ 111237 (growth zone), and YPM IZ 111240 (tentacle). B, *Physalia* are observed throughout the world, as shown by observations posted to inaturalist.org (red). Samples for genomic analysis (blue) were collected by an international collaboration of scientists. C, The first three principal components of genomic variation reveal five clusters labeled A, B1, B2, C1, and C2.

## Results

### Reference genome

We generated a new genome assembly for *Physalia* from a specimen collected in Texas, USA in 2017. This assembly, along with its alternate haplotype counterpart, has high contiguity (N50 of 10.4 and 4.6 megabases, see Table S1) and high `BUSCO` completeness scores (89.7% and 86.9%, Table S2). The length of the primary and alternate assemblies are 3.33 and 2.69 gigabases (Gb) respectively, and like other siphonophores (*31*), the *Physalia* genome is characterized by a substantial fraction of repeat sequences (∼65%). To assess genome size variation, we used a k-mer analysis to estimate the genome size of 11 specimens that were sequenced to an coverage >20x of a 3.3Gb genome. This analysis estimated that genome sizes vary between 1.5 and 2.0 Gb (Fig. S1), indicating a potentially inflated number of repeat sequences in the reference assembly (note however that these estimates are significantly smaller than previous estimates based on flow cytometry that estimated the size as 3.2 Gb (*30*)). To account for this in all downstream analyses we used only reads mapped to non-repeat regions of the primary assembly. In addition to the genome assembly, we also generated a new transcriptome using full-length cDNA generated with PacBio Iso-Seq on an additional specimen of *P. physalis*, collected in Florida in 2023. We tested the robustness of our results to reference assembly by repeating analyses over both the genome and transcriptome assemblies.

### Distinct clusters

To test hypotheses about *Physalia* diversity and population structure, our team collected >350 specimens, the majority of which were deposited at the Yale Peabody Museum (Fig 1, and see supplementary text). We sequenced the genomes of 133 samples, 123 of which were identified as high quality datasets, and performed a principal component analysis (PCA). Results show samples are divided into five clusters along the first two principal components of genomic variation (Fig. 1C, S2). These five clusters are labeled as A, B1, B2, C1, and C2, given the adjacency of the latter pairs to one another along principal components. We repeated the PCA including the ten sequenced samples found to be of moderate (rather than high) quality, and observed the same five clusters (Fig. S3). We also repeated the analysis mapping reads to the Iso-Seq transcriptome reference, and observed the same results (Fig. S4), indicating that population genomic studies similar to those presented here may not require reference genomes.

### Genomic differentiation

The geographic distribution of the five clusters shows that at least two were observed across multiple ocean basins (Fig. 2A): cluster B1 was found in the S. Atlantic, S. Indian, and S. and N. Pacific; cluster C1 was found on both sides of the S. Indian and S. Pacific oceans. By contrast, cluster A was observed only in the N. Atlantic, B2 on the northernmost sampling locations on both sides of the N. Pacific, and C2 only in New Zealand and Tasmania. We evaluated genomic differentiation by calculating the reciprocal fixation index (*Fst*), averaged across non-repeat windows. Average *Fst* values range from 0.29 between B1 and B2, to 0.64 between A and C1, suggesting little genetic exchange between any pair of clusters (Fig. 2B, see Fig. S5 for range across genomic windows). Estimates of nucleotide diversity, *pi*, indicate that cluster A has the lowest overall diversity and clusters B1 and C2 have the highest (Fig. S6), consistent with estimates of individual heterozygosity (Fig. S1B).

**Figure 2:**
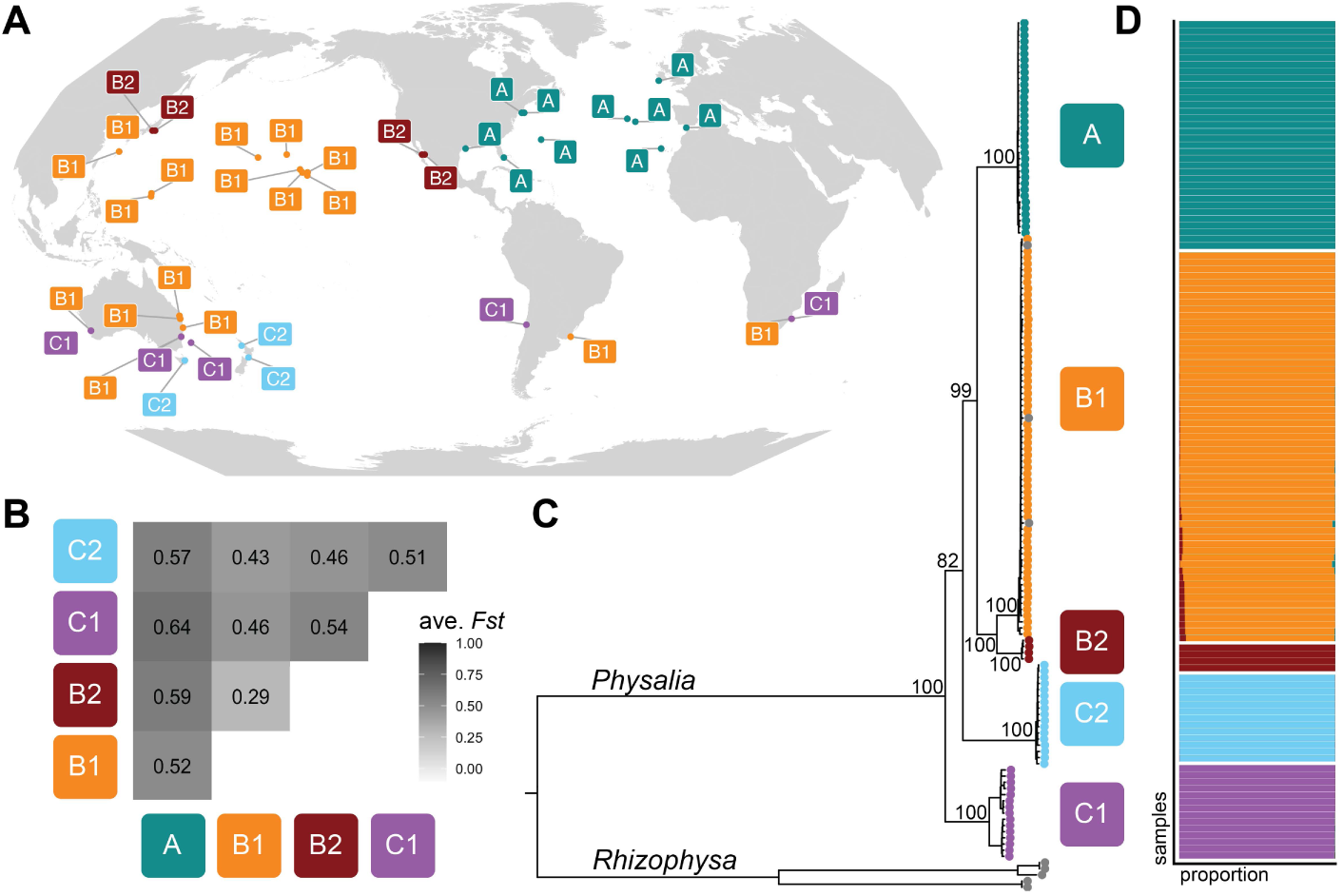
Multiple lines of evidence indicate reproductive isolation between lineages. A, The distribution of the five clusters from Fig. 1 shows some lineages span multiple ocean basins (e.g., B1, C1) and others are restricted to smaller areas (e.g., C2 observed in New Zealand and Tasmania). Labels indicate cluster present at collecting site. B, Reciprocal fixation index (*Fst*) averaged across non-repeat genomic windows indicate high levels of reproductive isolation between all lineages, with the weakest between B1 and B2. C, Phylogenetic analysis of 141 mitochondrial genomes shows reciprocal monophyly of lineages. Bootstrap values are shown at internal *Physalia* nodes. D, Shared ancestry analysis of 123 samples recovers five lineages with little evidence of mixture.

We tested the monophyly and phylogenetic relationships of these genomic clusters using two approaches. First, we assembled mitochondrial genomes for each sample and inferred a mitochondrial tree. For this analysis, we combined the mitogenomes generated in this study with all publicly available *Physalia* mitogenomes, and included all publicly available mitogenomes of the closely related genus *Rhizophysa* as an outgroup. The most likely tree shows clusters are monophyletic, with relatively little sequence variation within clusters (Fig. 2C). Clusters B1 and B2 were found to be sister to one another with high bootstrap support, and the clade of B1+B2 sister to cluster A. Support values were lower (bootstrap of 82) for the relationships at the base of the *Physalia* phylogeny.

Second, we estimated the phylogeny from a dataset of 800k high-quality SNPs, using the coalescent-based software `SVDQuartets`. A phylogeny of all specimens confirmed the reciprocal monophyly of the five lineages (Fig. S7). We examined the relationships between lineages by estimating a tree with individuals assigned to their respective clusters. Our results indicated a split between the clade (C1, C2), and the clade of (A and B1+B2) (Fig. S7C). Support values for both partitions in this unrooted tree showed unanimous support (bootstrap of 100).

We used a shared ancestry analysis to understand how genetic variants are partitioned across these lineages. The results favored five ancestry groups, corresponding to the five clusters above, and showed little evidence of mixture between groups (Fig. 2D, see Table S2 for D-statistics indicating no significant signatures of introgression). Repeating this analysis including the ten samples of moderate quality returned the same general results (Fig. S3), with the exception of three moderate-quality specimens of C2 that showed a minor proportion of mixture with C1 (Fig. S3). Repeating analyses using the reference transcriptome returned the same results (Fig. S4).

Several studies have generated data on individual genetic markers from *Physalia* (*17*, *32*– *35*). In order to place those data in the context of our findings, we inferred individual trees for four loci: mitochondrial CO1 and 16S, and nuclear ITS and 18S (Fig. S8-S11). We combined publicly available sequences from the National Center for Biotechnology Information (NCBI) with assembled marker sequences from our specimens, inferred using *in silico* PCR as implemented in our custom software `sharkmer`. This tool uses PCR primer sequences to seed a de Bruijn graph assembly of raw sequencing reads. These results furthered our understanding of *Physalia* diversity in the following ways: [1] a specimen reported from the Sargasso Sea (N. Atlantic) extended the predicted range of B1; [2] a specimen reported from Pakistan (N. Indian) extended the predicted range of B2; [3] using the internally transcribed spacer gene ITS, we were able to assign three clans, described in New Zealand (*17*), to clusters we describe here: clan 1 = cluster C2, clan 2 = cluster B1, clan 3 = cluster C1; however, using COI we found an incongruent result for the identity of clan 3. Without further information we cannot determine whether this result may be due to a potential exchange of mitochondrial sequences between clusters C1 and C2 in New Zealand.

### Morphology-based analysis

We tested for the evidence of distinct morphologies of *Physalia* by analyzing a dataset of images of *Physalia* uploaded to the citizen-science website inaturalist.org. While most of these images are of beached specimens, many aspects of the gross morphology are often preserved and identifiable. We scored the following characters (Fig. 3A): the height and length of the sail relative to the float; the color of the sail apex and the colony bodies (primarily gastrozooids); the arrangement of principal tentacles (defined as those with dense aggregations of tentilla); and the visible presence of a gap between the posterior and main zone of the colony. We scored characters on a dataset of 4,047 images, selected to include multiple images from all represented countries and time zones, with additional images scored for locations hypothesized to have increased diversity (e.g., New Zealand (*17*)). To ensure reproducibility of scoring, we had three independent observers score the same set of 100 images.

**Figure 3:**
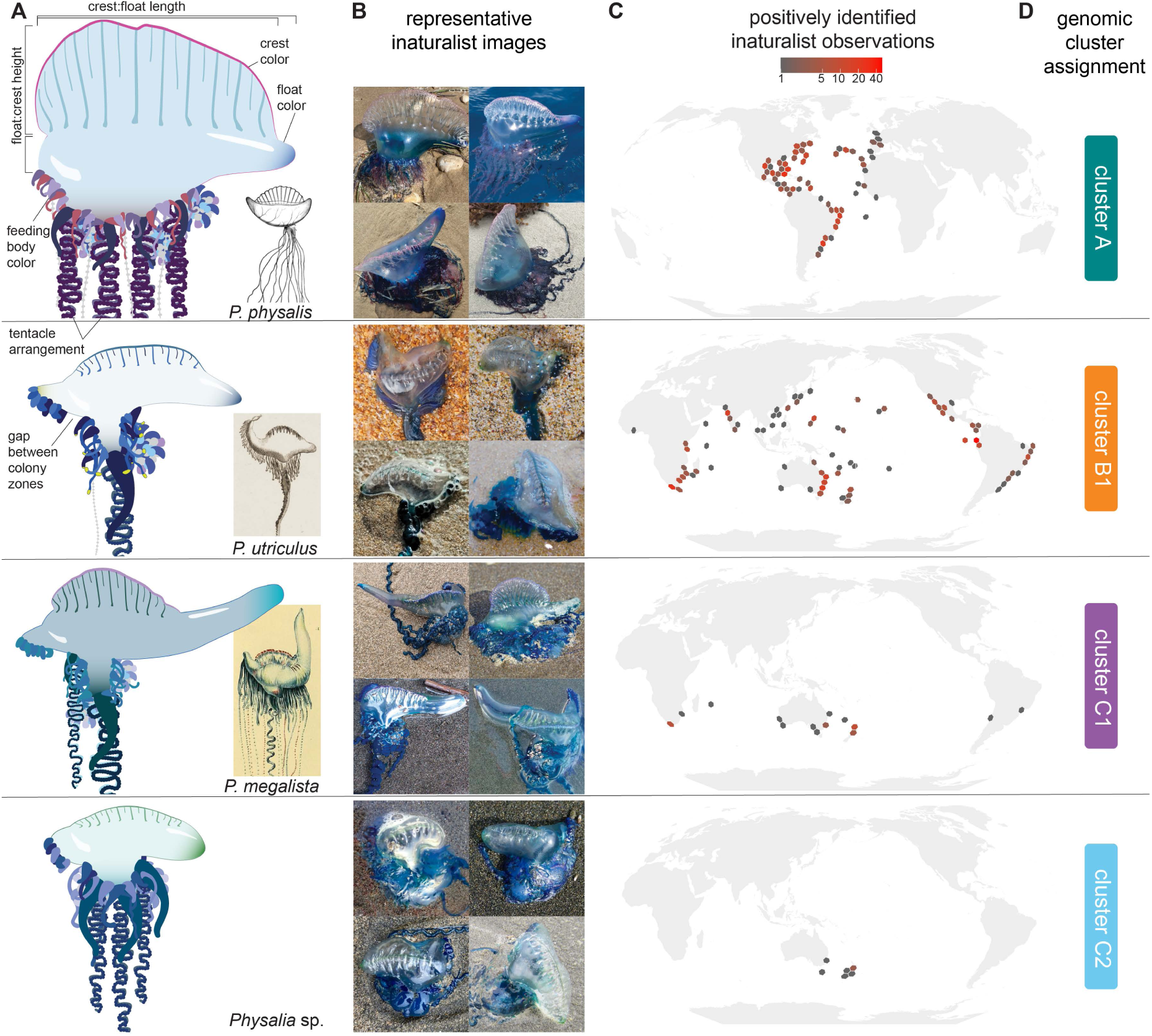
Distinct morphologies are detectable in citizen science images. A, Morphological traits such as aspects of size, color, and tentacle number were scored for thousands of images on inaturalist.org. From these, four morphologies were identified, three of which correspond to historically proposed species (*38*). B, Representative photos of each morphology from `iNaturalist`, image credits listed below. C, Ranges of positively identified `iNaturalist` records for each morphology, using a rule-based analysis of morphological traits. D, Morphologies were assigned to a genomic cluster by scoring the same traits of genomic specimens. Cluster B2 could not be definitively assigned due to lack of images and fixed material.

From these images, we identified four distinct morphologies (Fig. 3A-C). These were defined by describing a series of rules for positive identification based on suites of characters (Fig. S12), excluding images of poor quality or of specimens scored as having juvenile characteristics (e.g., globular float, few zooids). These rules constitute a strict definition for a high-confidence observation of each type; for example, images were positively identified as the *P. physalis* morphology if they had reddish feeding bodies, multiple major tentacles, and a sail that is as tall as the float and extends nearly to the anterior end. While individual specimens of *P. physalis* may deviate from these characters (e.g., if the sail is not raised), the rules were designed to minimize overlap between morphologies and allow for high-confidence identifications.

Three of the morphologies we identified are congruent with species proposed by scientists centuries ago (*16*). *P. physalis* was named by Linneaus in 1758 based on specimens from the Atlantic that had large sails and multiple major tentacles (Fig. 3A). *P. utriculus* was named by Gmelin (1788) (*36*), based on illustrations by La Martinière (1787) (*37*) of a Pacific specimen collected on the Lapérouse expedition that had a single major tentacle, yellow-tipped gastrozooids, and a flared posterior growth zone. *P. megalista* was named and illustrated by Lesueur and Petit (1807) (*38*) from specimens from the Southern Ocean that had an short sail and a sinuously postured float. Each of these species was synonymized with *P. physalis* in later centuries (*14*, *16*, *39*, *40*); our results indicate these synonymies to have been incorrect.

We linked these morphotypes to clusters identified through genome sequencing by analyzing the morphology of specimens we had analyzed genetically, using images taken upon collection, when available, as well as the morphology of fixed specimens (Fig. 3D, see supplementary text for specimen photos). Our results confirm that cluster A corresponds to *P. physalis*, B1 to *P. utriculus*, C1 to *P. megalista*, and C2 to *P.* sp. Cluster B2 could not be assigned given that no images of specimens were taken upon collection; analysis of the morphology of the single available fixed specimen suggested a general similarity to specimens of B1, *P. utriculus*.

Based on the assignment of morphotypes to clusters, we re-examined the distribution of the lineages using positively identified images (Fig. 3C). We found that cluster A, *P. physalis* was observed in the N. Atlantic, consistent with genomic findings, as well as the SW. Atlantic; B1, *P. utriculus* was found throughout the Pacific, Indian, as well as the SW. Atlantic and Gulf of Mexico; C1, *P. megalista* was found in the Southern edges of the Pacific, Indian, as well as the SW. Atlantic; and C2, *P. sp.* was found in New Zealand, Tasmania, as well as E. Australia.

### Geographic population structure

Given these animals can move with wind and currents, and combined with the evidence of distributions extended across ocean basins, we tested for evidence of long-distance dispersal and subpopulation structure by performing PCA on genomic variation within each of the four species: *P. physalis*, *P. utriculus*, *P. megalista*, and *P. sp.* C2. For *P. utriculus* we repeated this analysis both including and excluding cluster B2 (Fig. S13). Within species, samples are largely grouped by geographic region (Fig. 4), and not by date of collection (Fig. S14). The observation of a strong geographic signature, persistent even at sites with collection events over the span of multiple years, suggests that *Physalia* subpopulations largely stay in the same region over time. The extent of these geographic regions appears linked by major ocean currents; for example *P. physalis* specimens from Florida, Bermuda, and New England are highly similar, without substructure corresponding to collection sites, indicating these samples are part of a subpopulation aligned with the Gulf Stream current system (Fig. 4A, E).

**Figure 4:**
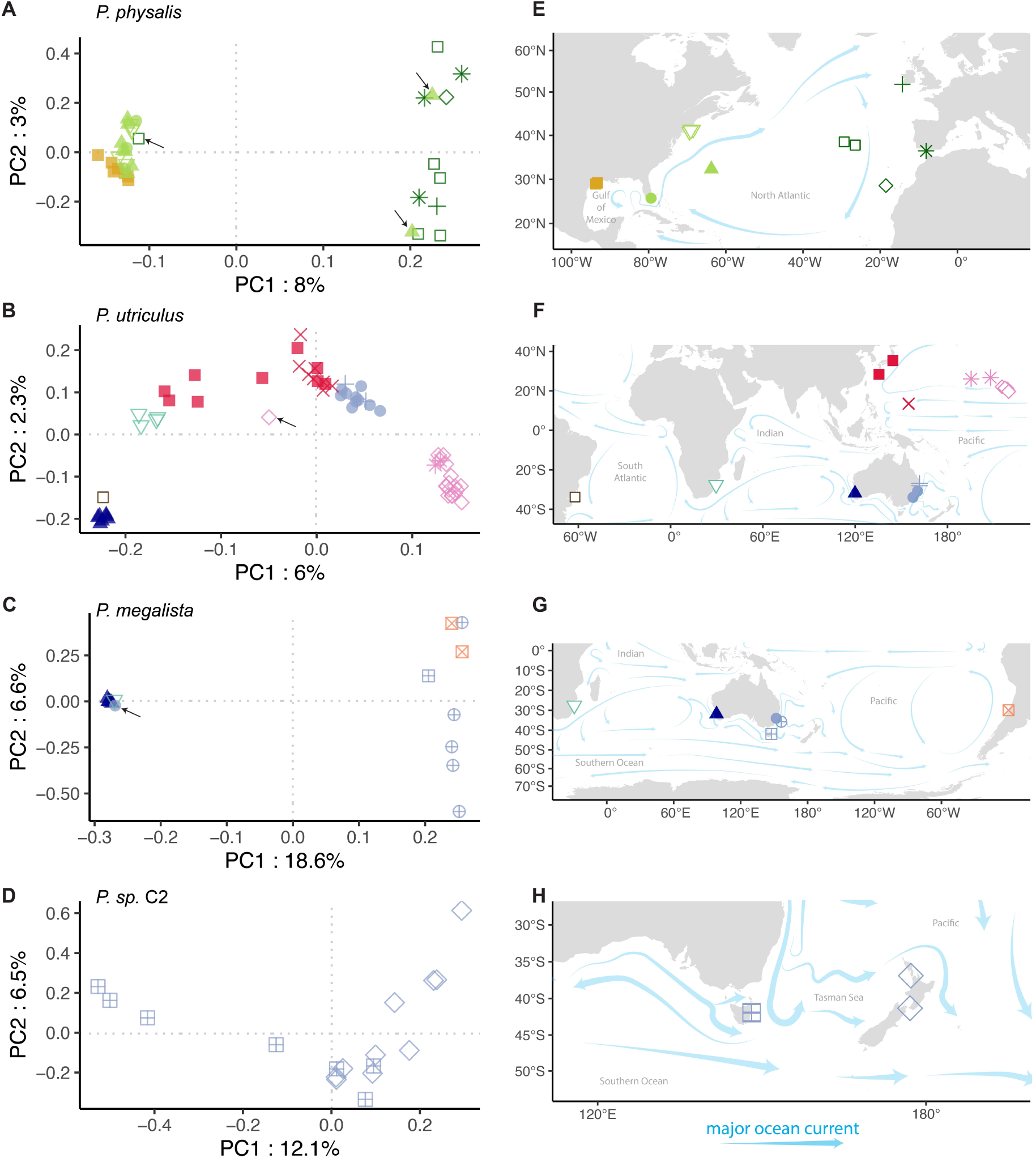
Principal component analyses within species show that subpopulations are largely defined by region. Exceptions to this pattern are marked with black arrows; these individuals suggest long-distance dispersal events across regions. A, *P. physalis*, B, *P. utriculus*, C, *P. megalista*, and D, *P.* sp. C2. Colors indicate regions of the ocean (e.g., Northwest Atlantic), and shapes indicate sampling location (e.g., Florida). E-H, Corresponding sampling locations and major ocean currents are shown, currents from National Oceanic and Atmospheric Administration (*42*).

We observed several individual exceptions to the pattern of persistent regional subpopulations within the dataset, indicating individuals can disperse over long distances (Fig. 4, S7A). In *P. physalis*, two of the samples collected in Bermuda showed an E. Atlantic subpopulation signature, and one sample in the Azores had a W. Atlantic signature, suggesting dispersal events in both directions across the N. Atlantic. In *P. utriculus*, one sample collected in Hawai’i showed a genomic signature associated with samples collected in Guam and Japan, suggesting an individual eastward dispersal event across the N. Pacific. In *P. megalista*, one sample collected in E. Australia had an Indian Ocean signature, suggesting dispersal across ocean basins.

We also tested the strength of differentiation between subpopulations within species. Using a k-means clustering analysis, we identified two subpopulations within *P. physalis*, three in *P. utriculus*, and two each in *P. megalista* and *P. sp.* C2 (Fig. S15). Lineage B2 was treated as a single cluster, given limited sampling. Genomic differentiation between subpopulations (calculated as average *Fst*) was small (<0.05), with the exception of the division within *P. megalista* (average *Fst* of 0.103, Fig. S16), a division also reflected in the mitochondrial and nuclear phylogenetic results (Fig. 2C, S7A), suggestive of a barrier to gene flow within *P. megalista*. We also examined genomic differentiation between specimens from different species that shared the same sampling region (Fig. S17), and confirmed that genomic differentiation is equivalent for groups of specimens regardless of co-occurence (e.g., *P. utriculus* and *P. megalista* that both occur in the SW. Pacific, average *Fst* value of 0.459).

## Discussion

This study, targeting an organism capable of long distance dispersal via ocean currents and winds, suggests that panmictic, cosmopolitan populations are the exception, and not the rule, for marine invertebrates. Our results show multiple lines of evidence that there are at least four species of *Physalia*, each composed of regionally endemic subpopulations. These lines of evidence include high genomic differentiation (measured as average reciprocal *Fst*), reciprocal monophyly of mitochondrial and nuclear phylogenies, clear morphological differentiation, and consistent mapping between genomic and morphological groupings. The four species we identify have distinct but overlapping ranges: *Physalia physalis* from the Atlantic; *P. utriculus*, present throughout the Pacific, Indian, and into the SW Atlantic; *P. megalista*, present in the southern portion of the Pacific, Indian, and Atlantic, and *P.* sp. C2, present in the Tasman Sea. We also find evidence to suggest a potential fifth species (cluster B2), but the absence of morphological data for sequenced specimens, combined with the relatively lower *Fst* value and phylogenetic proximity to *P. utriculus*, precludes its designation at this time.

Within species we observe genomic signatures endemic to specific regions, which are persistent over multiple years of sampling. In the case of *P. utriculus* in Hawai’i, we collected reproductively mature adults (e.g., YPM IZ 110777), juveniles collected before surfacing (e.g. YPM IZ 110881, YPM IZ 110882, YPM IZ 110883, collected at ∼6 meters depth), and a range of sizes in between (see supplementary text), and observed that they shared the same subpopulation signature, suggesting reproduction happens *in situ*. Regional genomic signatures are robust despite our observation of five individual specimens with incongruent genomic and geographical signatures that suggest cross-regional dispersal events. Subpopulation boundaries appear to be defined by patterns of winds and currents, as demonstrated by the close genetic affinity of samples collected at sites adjacent to the Gulf Stream (Texas, Florida, Bermuda, and New England).

The southern hemisphere, and in particular the southwest regions of ocean basins, consistently represent centers of *Physalia* diversity: three species are found in the SW. Pacific (*P. utriculus*, *P. megalista*, and *P.* sp. C2), two species in the S. Indian Ocean (*P. utriculus* and *P. megalista*), and three species in the SW. Atlantic (*P. physalis*, *P. utriculus*, and *P. megalista*). In no case, do we observe evidence of gene flow between species at the sites of range overlap (Fig. S17). Furthermore, the phylogenetic relationships between species suggest that diversification may have originated from the Pacific Ocean. The most recent species division is between *P. physalis* and *P. utriculus*, potentially as the ancestral population of these lineages came to occupy the Atlantic Ocean. In addition, we observe moderate genomic differentiation between clusters B1 and B2 in the N. Pacific (ave. *Fst* of 0.29, Fig. 2), and between subpopulations of *P. megalista* in the Pacific and Indian Oceans (ave. *Fst* of 0.12, Fig. S16), both suggesting population structure due to potential ecological or geographic divides within species.

This study builds on the work and observations of sailors, swimmers, and scientists over the course of centuries. As early as the 18th century, hypotheses about multiple species emerged, based on reports from global voyages (*16*). Among these are three of the species we observed, *P. physalis*, *utriculus*, and *megalista*. These species were not “cryptic”; they were proposed, debated, and ultimately rejected over the course of 250 years. Our results vindicate their original descriptions, showing clear and strong support for distinct species matching the original illustrations. The central challenge faced by taxonomists in past centuries was that there was no way to simultaneously observe live or recently beached *Physalia* across its huge range, and key characteristics like posture, color, and behavior are lost during fixation. These results underscore the power of participatory science and social media to provide an unprecedented lens on biodiversity.

Conflicting expectations and observations of the number of planktonic species have spawned multiple discourses in the literature (e.g., “the paradox of the plankton” (*43*) and its companion, “the inverted paradox of the plankton” (*44*)). Here we demonstrate that, in the case of *Physalia*, there is more diversity than previously assumed (four species instead of one), and that the open ocean ecosystem might indeed have high evolutionary potential (*4*). Across the open ocean we observe substantial geographic partitioning of genetic variation, evidence for reproductive isolation events that have resulted in strong barriers to reproduction, as well as events that suggest isolation may be currently underway (e.g. clusters B1 and B2), and in the case of *P. sp.* C2, we report a previously undescribed species that represents a single-sea endemic. Future research into the physical, environmental, and biological processes that generate and maintain this genetic variation will be crucial in recalibrating our expectations towards open-ocean biodiversity.

## Supporting information

supp. specimen information

## Acknowledgments

We thank Adachi Aya, Stephen Chiswell, Lynne Christianson, Gary Cobb, Andy Coetzee, Joey DiBatista, Steve Dunn, Leanne Elder, Jose Fernández-Simón, Luke Finley, Steve Gerber, Rosemary Hill, Kazuhisa Hori, Takato Izumi, Michiyo Kawabata, Itaru Kobayashi, Hisanori Kohtsuka, Izumi Komori, Hiromi Kurihara, Yukio Kurihara, Nate Lo, Juan Moles, Kirrily Moore, Natasha Picciani, Nicola Puharich, Fae Spafford, John Stimson, Hiroyuki Tachikawa, Nikolai Tatarnic, Yi-Kai Tea, Kosuke Uchiwa, Diego Vazquez Alcaide, Kensuke Yanagi, the Miura Fisher Cooperative Association, and the East Matunuck State Beach lifeguards for additional support in collecting specimens. We thank Lawrence Gall, Gisella Caccone, and the Caccone laboratory for early feedback. We thank the Western Australian Museum, Tasmanian Museum and Art Gallery, and the Field Museum of Natural History for loaning specimens. We also thank the Bingham Oceanographic Collection Endowment, the Yale Center for Genome Analysis, and the Keck Microarray Shared Resource at Yale University for providing necessary sequencing services.

## Image credits

Images reproduced in Figure 3B were retrieved from https://inaturalist.org and are all made available under Creative Commons licence CC-0 or CC-BY. They are credited as follows, left to right and top to bottom:

1. Eileen Mattei, Texas, USA. CC-0. https://inaturalist-open-data.s3.amazonaws.com/photos/271981781/original.jpg
2. Seth Wollney, Atlantic Ocean. CC-BY. https://inaturalist-open-data.s3.amazonaws.com/photos/314857468/original.jpg
3. Amanda Chase, Texas, USA. CC-BY. https://inaturalist-open-data.s3.amazonaws.com/photos/4323195/original.jpeg
4. Lucian Vazquez, Matanzas, Cuba. CC-BY. https://inaturalist-open-data.s3.amazonaws.com/photos/249736410/original.jpg
5. Tim (user twan3253), New South Wales, Australia. CC-BY. https://inaturalist-open-data.s3.amazonaws.com/photos/116503028/original.jpeg
6. William Stephens. San Cristobal, Ecuador. CC-BY. https://inaturalist-open-data.s3.amazonaws.com/photos/64365445/original.jpg
7. Andrew Gillespie, Eastern Cape, South Africa. CC-BY. https://inaturalist-open-data.s3.amazonaws.com/photos/349817115/original.jpeg
8. Manuel R. Popp, Eastern Cape, South Africa. CC-BY. https://inaturalist-open-data.s3.amazonaws.com/photos/180673664/original.jpeg
9. Hazel Valerie, Northland, New Zealand. CC-BY. https://inaturalist-open-data.s3.amazonaws.com/photos/145998383/original.jpeg
10. Jacqui Geux, Auckland, New Zealand. CC-BY. https://inaturalist-open-data.s3.amazonaws.com/photos/3560793/original.JPG
11. Emily Roberts, Taranaki, New Zealand. CC-BY. https://inaturalist-open-data.s3.amazonaws.com/photos/205384503/original.jpeg
12. Arnim LIttek, Manawatu-Wanganui, New Zealand. CC-BY. https://inaturalist-open-data.s3.amazonaws.com/photos/227840858/original.jpeg
13. Arnim LIttek, Manawatu-Wanganui, New Zealand. CC-BY. https://inaturalist-open-data.s3.amazonaws.com/photos/72081497/original.jpeg
14. Jacqui Geux, Auckland, New Zealand. CC-BY. https://inaturalist-open-data.s3.amazonaws.com/photos/65125848/original.jpeg
15. Roderic Page, Auckland, New Zealand. CC-BY. https://inaturalist-open-data.s3.amazonaws.com/photos/340905787/original.jpg
16. Arnim LIttek, Manawatu-Wanganui, New Zealand. CC-BY. https://inaturalist-open-data.s3.amazonaws.com/photos/36406644/original.jpeg

## Author contributions

1. SHC-NA, KBD, MKM, CM, EALW-CWD contributed to conceptualization
2. SHC-NA, CWD contributed to data curation and validation
3. SHC-RBA, CWD contributed to formal analysis
4. EALW, CWD contributed to funding acquisition
5. SHC-NA, DAR, OB, KBD, AGR, ZRL, AdM, CM, CWD contributed to investigation (e.g., DNA extraction, sequencing)
6. SHC, CWD contributed to methodology
7. SHC-BIMMM, AdM-BMD, CWD contributed resources (e.g., specimens)
8. SHC-RBA, CWD contributed to software development
9. CS-CWD contributed to project supervision
10. SHC-RBA, SHDH, JMC-CWD contributed to visualization
11. SHC-CJA, ADS, SHDH, RRH, MKM, KAP, AS, EALW-CWD contributed to writing the original draft
12. SHC-CWD (all authors) contributed to reviewing and editing the final draft

## Data and materials availability

Sequence data for genome assembly (PacBio reads and 10X Genomics Chromium linked-reads) are available at the National Center for Biotechnology Information (NCBI) Bioproject PRJNA735958, and the principal and haplotype assembly sequences are available at NCBI Bioproject PRJNA1040906. Full-length, non-chimeric PacBio Iso-Seq RNA data are available at BioProject PRJNA1126252. Illumina sequence data for specimens intended for population genomic analysis are available at Bioproject PRJNA1092115. Sequence alignment and phylogenetic tree files, along with all code used to analyze data to reproduce the figures shown here (Snakemake workflows and Rmarkdown files) are available at https://github.com/shchurch/Physalia_population_genomics, commit 7a2a4ac. Specimens collected for this study are deposited at the Yale Peabody Museum, with the exception of specimens deposited at the New Zealand National Institute for Water and Atmospheric Research (NIWA) Invertebrate Collection, and those loaned by the Western Australian Museum, Tasmanian Museum and Art Gallery, and the Field Museum of Natural History. Specimen catalog numbers listed in supplementary text: specimen collection information.

## Materials and Methods

### Sample collection and DNA extraction

*Physalia* specimens were collected by a global collaboration of scientists. Full sampling details and required permit information can be found in the supplementary text: specimen collection information. The majority of specimens were collected after they washed ashore, using appropriate safety protocols to avoid stings. A few specimens were collected directly from the water, either sampling from a boat or while diving (for juvenile specimens that hadn’t yet surfaced). Specimens were preserved in >70% ethanol, when available, and stored at room temperature, with the exception of two samples collected from Hawai’i that were stored in DNAshield, and several samples from the Eastern United States that were flash-frozen.

When possible, whole specimens were collected and shipped to the Yale Peabody Museum, Invertebrate Zoology Division (YPM IZ). All sequenced specimens were photographed, and images are shown in the supplementary text. Additional specimens were loaned from the Western Australia Museum, the Tasmanian Museum and Art Gallery, and the Field Museum in Chicago.

High molecular weight DNA for genome sequencing and assembly was extracted from a flashfrozen specimen, YPM IZ 110876, collected in Texas, USA in 2017. Extractions were performed following the protocol described by Chen and Dellaporta, 1994 (*45*), with modifications. In this protocol, tissue was homogenized under liquid nitrogen and extracted with 5 mL of a urea-based extraction buffer for 15 minutes at 65 degrees C. Three 25:24:1 phenol:chloroform:isoamylalcohol (P:C:I) extractions were performed, each allowed to rock for 5 minutes before centrifugation. The P:C:I extractions were followed up with extraction with one volume of chloroform prior to precipitation with isopropanol. Extracted DNA and RNA was resuspended in 100 *μ*l Tris-EDTA buffer, analyzed by gel electrophoresis, then brought to a volume of 400 *μ*L and subjected to RNase treatment with 3 *μ*L RNase I and 2 *μ*L of RNAse Cocktail Enzyme Mix for 60 minutes at 37 degrees C. The RNA-free DNA was then brought to 500 *μ*l with 5M NaCl and extracted with 400 *μ*l P:C:I. The aqueous phase was removed and 400 *μ*L of 5M NaCl, 500 mM EDTA, 10 mM Tris was added to the P:C:I for back extraction. The back-extracted aqueous phase was combined with the first aqueous phase and 0.3 volumes of 100% ethanol was added to precipitate polysaccharides, pelleted at 17,000 x G. DNA was precipitated with 1.7 volumes of 100% ethanol, pelleted, and washed with 70% ethanol and resuspended in Tris-EDTA. The purified DNA was examined by pulse-field gel electrophoresis and showed a strong band at >98kb.

RNA for transcriptome sequencing was extracted from a flash-frozen specimen, YPM IZ 110436, collected in Florida, USA in 2023. Tissue was homogenized using a mortar and pestle chilled to −80 degrees C, and RNA was extracted using the RNAqueous Total RNA Isolation Kit following manufacturers instructions and including a lithium-chloride precipitation step. RNA was processed for library preparation and PacBio Iso-Seq sequencing by the Keck Microarray Shared Resource at Yale University. Delivered reads were clustered with the PacBio isoseq cluster2 command, version 1.0.1. These were then further deduplicated with treeinform (*46*) as implemented in the code available at https://github.com/dunnlab/isoseq, commit 7a2a4ac. This tool implements a phylogenetically informed refinement of the transcriptome to remove species-specific variants, by building gene trees from the target transcriptome (here *Physalia*) and gene predictions for related species (here 23 species, including 10 cnidarians). Clades with short total branch length and that contain only sequences from the target species are collapsed to the longest sequence.

For samples intended for population genomic analysis, tentacle pieces were dissected from whole specimens and stored in 95% ethanol prior to DNA extraction for genome sequencing. DNA extractions were performed using the EZNA Mollusk kit following manufacturers instructions and an overnight digestion, with the exception of several samples from Japan, Guam, and Texas that were extracted using the urea-based phenol-chloroform protocol described above, as well as one sample from the Gulf of California, extracted at Monterey Bay Aquarium Research Institute with the DNeasy DNA Blood and Tissue kit (Qiagen), following manufacturer’s instructions. Whole genome DNA was processed for library preparation and sequencing by the Yale Center for Genome Analysis.

### Genome assembly

Eight Single-Molecule Real-Time (SMRT) sequencing cells of PacBio HiFi data were assembled with `canu`, v. 2.2 (`-pacbio-hifi` option) (*47* –*49*), with the estimated genome size parameter set to three gigabases. HiFi reads were mapped to this assembly with `minimap2`, v. 2.22-r1101 (*50*), to determine the appropriate cutoffs for purging duplicated contigs. These were removed using `purge_haplotigs`, v. 1.1.2 (low, medium, and high cutoffs set at 5x, 40x, and 200x respectively) (*51*), and overlapping contig ends were clipped with the same program. The parameters for `purge_haplotigs` were modified to avoid memory limitation (-I was set to 1G, -p was dropped, and -N was set to 1000).

A foreign contamination screen (FCS, via the National Center for Biotechnology Information, NCBI) was performed on both the purged and haplotype assemblies, using the tool provided for GenBank submissions which detected and removed one adapter sequence. We used the tool `LongStitch`, v. v1.0.4 (*52*), to scaffold both the purged (primary) and haplotype (alternate) assemblies. Scaffolding was performed first using the eight HiFi cells used for assembly and the `ntLink-arks` functionality, and then using a dataset of 225 gigabases of linked-read data sequenced with 10XGenomics Chromium sequencing, interleaved with `LongRanger` (provided by 10X Genomics). The FCS was repeated on this assembly and detected no further foreign contaminants.

Repeat regions were detected and masked with `RepeatModeler` and `RepeatMasker`, v. 4.1.5 (*53*), to build a general feature format (gff) file, used to exclude repeats from downstream analyses. `BUSCO`, v 5.4.4 (*54*), and `BBMap stats.sh` were used to evaluate final assemblies. Genome assembly is made publicly available at NCBI, BioProject number PRJNA1040906.

### Genome mapping

Paired-end genome sequencing targeting a read length of 150 base pairs was performed for 145 libraries using an Illumina NovaSeq at the Yale Center for Genome Analysis. Full details on quality control, mapping statistics, and final library parameters are available in the GitHub document https://github.com/shchurch/Physalia_population_genomics/manuscript_files/quality_control.html. Briefly, sequencing depth range varied across samples from a target of 10-60x genome size. These 145 samples included two replicate libraries, generated from repeated DNA extractions from the same specimens. In addition, from sequenced libraries we generated two technical replicates by randomly splitting read files. These replicates were used to evaluate reproducibility and were excluded from the main analyses presented in this work.

Overall sequence quality (e.g. GC content, adapter content) was evaluated using `FastQC`, v. 0.11.9. Reads were trimmed for Illumina adapters using `Trimmomatic`, v. 0.39 (*55*). Potential human, bacterial, and viral DNA contamination was evaluated using Kraken2, v. 2.1.2 (*56*), standard database. Additional cross-species contamination was evaluated using *in silico* PCR of the ribosomal 18S gene from genomic reads, and comparing results to publicly available datasets with a basic local alignment search tool, `BLAST`. Potential kinship or cross-contamination between *Physalia* samples was evaluated by calculating the kinship-based inference for genomes (KING-robust) relatedness score on reads mapped to the assembled genome using `PLINK2`, v. 2.00a5LM (*57*), calculated only using SNPs within Hardy-Weinberg equilibrium (p-value <1e-7), and excluding those with missing alleles >0.1 or a minor allele frequency >0.01).

Based on the results of the quality control analyses, six samples were identified as contaminated and an additional four samples were identified as replicated sampling events from a single specimen (e.g. multiple tentacle tips taken from the same animal in the field). These samples were excluded from downstream analyses, such that the final dataset, excluding technical and biological replicates, consisted of 133 samples. Of those, 123 were marked as high quality based on overall sequencing depth, read quality, and proportion of missing sites. Analyses were performed on a strict dataset of only high-quality samples, and repeated on the full dataset of high- and moderate-quality samples.

Reads were mapped to the reference genome using `BWA`, v. 0.7.17-r1188 (*58*). Mapped reads were sorted, deduplicated, and indexed using `picard`, v. 2.25.6. Alleles were called using `BCFtools`, v. 1.16, `mpileup` (*59*). To test the robustness of downstream analyses to reference assembly, reads were mapped to the independent transcriptome assembly, using only R1 reads as single-end data.

### Genome size estimates

We estimated the genome size for 19 specimens sequenced to >20x coverage, estimated against a genome size of 3.3 Gb. Genome size, repeat content, and heterozygosity were estimated using GenomeScope, v. 2.0 (*60*) and jellyfish, v. 2.2.3 (*61*). GenomeScope was run on the combined set of R1 and R2 reads, trimmed to remove Illumina adapters as described above. Size estimates were discarded for eight samples with a maximum model fit <95% or when the fitted model failed to follow the curve coverage histogram.

### Phylogenetics

Mitochondrial genomes were assembled from a subset of ten million trimmed reads for each sample, using the software `GetOrganelle`, v. 1.7.7.0 (*62*), using the `animal_mt` database and default parameters. `GetOrganelle` failed to circularize the assemblies, in line with the expected linear mitochondrial genomes in siphonophores (*31*); the resulting top path assembly was used as the final linear genome. Assembled sequences were combined with publicly available mitochondrial assemblies for *Physalia* and their outgroup *Rhizophysa* from NCBI, accession numbers: OQ957220, KT809328, LN901209, KT809335, NC_080942, NC_080941, OQ957206, OQ957199. Mitochondrial genomes were aligned using `MAFFT`, v. 7.505, `--adjustdirectionaccurately` option (*63*). A mitochondrial phylogeny was inferred using `IQtree2`, v. 2.2.6 (*64*), model autoselected (*65*) and 1,000 ultrafast bootstraps (*66*), with *Rhizophysa* selected as the outgroup.

Individual marker sequences were assembled from raw reads using *in silico* PCR as implemented in `sharkmer` (available at https://github.com/caseywdunn/sharkmer, commit `c43cfc2`). Four markers were selected to infer individual gene trees: mitochondrial cytochrome oxidase I (CO1), mitochondrial large ribosomal subunit 16S, nuclear ribosomal internal transcribed spacer (ITS), as well as small nuclear ribosomal subunit 18S. These markers were combined with all publicly available *Physalia* and *Rhizophysa* sequences for the same genes, from NCBI. Sequences were aligned with `MAFFT`, and gene trees inferred with `IQtree2`, as described above.

A phylogeny of single nucleotide polymorphisms (SNPs) was assembled using `SVDquartets`, as implemented in PAUP*, v. 4.0a (*67*). SNPs were selected based on the following filters: minimum Phred quality of 40, minimum and maximum depth of 2x and 99x respectively, maximum proportion of missing data of 25%, minimum distance between SNPs set to 100 base pairs, excluding sites with only alternative alleles called, and only selecting bi-allelic SNPs. The final dataset contained 839,510 SNPs. `SVDquartets` was used to infer a phylogeny of all specimens without population-level information, and a phylogeny with specimens assigned to populations based on results of the principal component and shared ancestry analyses. For the latter, support was evaluated using 100 bootstraps.

### Principal components analysis

Principal component analysis (PCA) was performed on estimated genotype likelihoods, calculated using `ANGSD`, v. 0.935 (*68*), on reads mapped to a random sample of 100,000 non-repeat genomic regions, each larger than 1,000 base pairs. Sites were included based on the following filters: p-value of variability below 1e-6, minimum Phred quality score of 40, minimum and maximum depth of 2x and 99x respectively, and present in a minimum of 92 individuals (75% of 123 samples). PCA and ancestry analyses were performed using `PCANGSD`, v. 1.21 (*69*), `-admix-alpha` set to 50 and allowing the software to choose the optimal number of components.

PCA and shared ancestry analyses were repeated on the full subset of samples, using a minimum of 100 samples (75% of 133 samples), as well as with reads mapped to the reference transcriptome. PCA was also performed within each lineage detected in this study. Subpopulations were classified using k-means clustering of the resultant covariance matrices, with the optimal number of clusters chosen using an elbow plot of eigenvalues.

### Population statistics

Populations genomic statistics (*pi*, *Dxy*, and *Fst*) were calculated using `pixy`, v. 1.2.7 (*70*), on a dataset of alleles filtered with the following metrics: minimum Phred quality score of 40, minimum and maximum depth of 2x and 99x respectively, maximum missingness of 25%. Statistics were calculated on a random sample of 100,000 non-repeat genomic regions, each larger than 1,000 base pairs, and summary statistics were averaged over these regions. Statistics were calculated between lineages as assigned using PCA and ancestry analyses; between subpopulations, as defined using PCA and ancestry within species; and between lineage + sampling location combinations.

D-statistics were calculated using `Dsuite`, v. 0.5 r57 (*71*), the `Dquartets` function on the same dataset of filtered alleles. Populations were defined using the results of the shared ancestry analysis on 133 high and moderate quality samples. Significant signatures of introgression were defined as having a Z-score >2 and a p-value <0.05.

### Morphological scoring of images

ID numbers for ∼11,000 research-grade photos of *Physalia* were downloaded from inaturalist.org in October, 2023. Of these, a subset of 4,047 images were scored, selected to include multiple images from all represented countries and time zones, as well as to maximize representation in areas hypothesized to have increased diversity (specifically New Zealand, South Africa, and Brazil). Images were categorized based on quality and perspective on the animal (e.g., ventral, dorsal, or lateral), and were scored for the following traits:

- sail height, binned into four categories: as tall as float, >1/3 the height of float, <1/3 the height of float, or flush with float / no visible height
- length of float anterior to the end of the sail, binned as <1/4 sail length, >1/4 and <3/4 sail length, and >3/4 sail length
- presence of pink or purple coloration on the sail
- presence of yellow or reddish coloration on gastrozooids
- clear, glassy float coloration
- arrangement of principal fishing tentacles (defined as having tentilla tightly packed), categorized as having one central tentacle, two central tentacles, or many
- presence of a gap between the central (main) and posterior colony zone of zooids
- juvenile morphology, defined as having a globular float with one or no major tentacles, no sail height, and few zooids.

Each trait was only scored when visible, therefore absence of a score is not evidence of trait absence. Images were scored in batches by three different researchers (SHC, RBA, and NA). To ensure consistency, researchers independently scored the same set of 100 randomly sampled photos, and compared results to bring qualitative assignments into alignment. Images classified as being of poor quality, taken from a ventral perspective, or of a juvenile specimen as defined above, were excluded from downstream analyses.

Four morphological types were identified from scored images in combination with descriptions and diagrams of historically hypothesized species. Rules for assigning images to one of these four morphologies were established based on combinations of characters, see Fig. S12. Given the potential plasticity of the traits in question (e.g., color, size), no single trait was considered diagnostic. Genomic clusters were associated with these morphologies by scoring the same traits on the specimens processed for genomic analyses.

When image assignments extended the known range of a genomically defined lineage, these images were independently rescored by two researchers. If there was any discrepancy in the resulting scores for a trait relative to the morphological assignment, the image was excluded from the rule-based analysis.

## References

1. S. Van der Spoel, The basis for boundaries in pelagic biogeography. Progress in Oceanography 34, 121–133 (1994).

2. M. V. Angel, Biodiversity of the pelagic ocean. Conservation Biology 7, 760–772 (1993).

3. R. D. Norris, Pelagic species diversity, biogeography, and evolution. Paleobiology 26, 236–258 (2000).

4. K. T. Peijnenburg, E. Goetze, High evolutionary potential of marine zooplankton. Ecology and Evolution 3, 2765–2781 (2013).

5. B. W. Bowen, M. R. Gaither, J. D. DiBattista, M. Iacchei, K. R. Andrews, W. S. Grant, R. J. Toonen, J. C. Briggs, Comparative phylogeography of the ocean planet. Proceedings of the National Academy of Sciences 113, 7962–7969 (2016).

6. S. B. Johnson, J. R. Winnikoff, D. T. Schultz, L. M. Christianson, W. L. Patry, C. E. Mills, S. H. Haddock, Speciation of pelagic zooplankton: Invisible boundaries can drive isolation of oceanic ctenophores. Frontiers in Genetics 13, 970314 (2022).

7. S. H. Haddock, A golden age of gelata: Past and future research on planktonic ctenophores and cnidarians. Hydrobiologia 530, 549–556 (2004).

8. J. B. Pfaller, A. C. Payton, K. A. Bjorndal, A. B. Bolten, S. F. McDaniel, Hitchhiking the high seas: Global genomics of rafting crabs. Ecology and Evolution 9, 957–974 (2019).

9. R. R. Helm, The mysterious ecosystem at the ocean’s surface. PLOS Biology 19, e3001046 (2021).

10. C. J. Anthony, B. Bentlage, R. R. Helm, Animal evolution at the ocean’s water-air interface. Current Biology 34, 196–203 (2024).

11. F. Chong, M. Spencer, N. Maximenko, J. Hafner, A. C. McWhirter, R. R. Helm, High concentrations of floating neustonic life in the plastic-rich north pacific garbage patch. PLOS Biology 21, e3001646 (2023).

12. Z. Long, Begin ocean garbage cleanup immediately. Science 381, 612–613 (2023).

13. I. T. Wysocki, M. Wang, C. Morales-Caselles, L. C. Woodall, K. Syberg, B. C. Almroth, M. Fernandez, L. Monclús, S. P. Wilson, M. Warren, D. Knoblauch, R. R. Helm, Plastics treaty text must center ecosystems. Science 382, 525–526 (2023).

14. A. K. Totton, Studies on *Physalia physalis* (L.). Pt. 1 Natural history and morphology. Discovery Reports 30, 301–368 (1960).

15. J. Bardi, A. C. Marques, Taxonomic redescription of the Portuguese man-of-war, *Physalia physalis* (Cnidaria, Hydrozoa, Siphonophorae, Cystonectae) from Brazil. Iheringia. Série Zoologia 97, 425–433 (2007).

16. P. Pugh, A history of the sub-order Cystonectae (Hydrozoa: Siphonophorae). Zootaxa 4669, 1–91 (2019).

17. D. Pontin, R. Cruickshank, Molecular phylogenetics of the genus *Physalia* (Cnidaria: Siphonophora) in New Zealand coastal waters reveals cryptic diversity. Hydrobiologia 686, 91–105 (2012).

18. M. J. Costello, P. Tsai, P. S. Wong, A. K. L. Cheung, Z. Basher, C. Chaudhary, Marine biogeographic realms and species endemicity. Nature Communications 8, 1057 (2017).

19. C. Munro, Z. Vue, R. R. Behringer, C. W. Dunn, Morphology and development of the Portuguese man of war, *Physalia physalis*. Scientific Reports 9, 15522 (2019).

20. Y. K. Okada, Développement post-embryonnaire de la Physalie Pacifique. Memoirs of the College of Science, Kyoto Imperial University. Series B 8, 1–26 (1932).

21. J. E. Purcell, Predation on fish larvae by *Physalia physalis*, the Portuguese man of war. Marine Ecology Progress Series. Oldendorf 19, 189–191 (1984).

22. A. Damian-Serrano, S. H. Haddock, C. W. Dunn, The evolution of siphonophore tentilla for specialized prey capture in the open ocean. Proceedings of the National Academy of Sciences 118, e2005063118 (2021).

23. E. Dodge Wangersky, C. E. Lane, Interaction between the plasma of the loggerhead turtle and toxin of the Portuguese man-of-war. Nature 185, 330–331 (1960).

24. C. J. Anthony, Beachside banquet: Ants’ appetite for shipwrecked siphonophores. Food Webs 38, e00332 (2024).

25. L. Prieto, D. Macías, A. Peliz, J. Ruiz, Portuguese Man-of-War (*Physalia physalis*) in the Mediterranean: A permanent invasion or a casual appearance? Scientific Reports 5, 11545 (2015).

26. J. L. Headlam, K. Lyons, J. Kenny, E. S. Lenihan, D. T. G. Quigley, W. Helps, M. M. Dugon, T. K. Doyle, Insights on the origin and drift trajectories of Portuguese man of war (*Physalia physalis*) over the Celtic Sea shelf area. Estuarine, Coastal and Shelf Science 246, 107033 (2020).

27. D. R. Pontin, M. J. Watts, S. P. Worner, “Using multi-layer perceptrons to predict the presence of jellyfish of the genus *Physalia* at New Zealand beaches” in 2008 IEEE International Joint Conference on Neural Networks (IEEE World Congress on Computational Intelligence) (IEEE, 2008), pp. 1170–1175.

28. A. Canepa, J. E. Purcell, P. Córdova, M. Fernández, S. Palma, Massive strandings of pleustonic Portuguese Man-of-War (*Physalia physalis*) related to ENSO events along the southeastern Pacific Ocean. Latin American Journal of Aquatic Research 48, 806– 817 (2020).

29. N. Bourg, A. Schaeffer, P. Cetina-Heredia, J. C. Lawes, D. Lee, Driving the blue fleet: Temporal variability and drivers behind bluebottle (*Physalia physalis*) beachings off Sydney, Australia. PLOS One 17, e0265593 (2022).

30. K. Adachi, H. Miyake, T. Kuramochi, K. Mizusawa, S. Okumura, Genome size distribution in phylum Cnidaria. Fisheries Science 83, 107–112 (2017).

31. N. Ahuja, X. Cao, D. T. Schultz, N. Picciani, A. Lord, S. Shao, K. Jia, D. R. Burdick, S. H. Haddock, Y. Li, others, Giants among Cnidaria: Large nuclear genomes and rearranged mitochondrial genomes in siphonophores. Genome Biology and Evolution 16, evae048 (2024).

32. A. Collins, Phylogeny of medusozoa and the evolution of cnidarian life cycles. Journal of Evolutionary Biology 15, 418–432 (2002).

33. A. G. Collins, S. Winkelmann, H. Hadrys, B. Schierwater, Phylogeny of Capitata and Corynidae (Cnidaria, Hydrozoa) in light of mitochondrial 16S rDNA data. Zoologica Scripta 34, 91–99 (2005).

34. C. W. Dunn, P. R. Pugh, S. H. Haddock, Molecular phylogenetics of the Siphonophora (Cnidaria), with implications for the evolution of functional specialization. Systematic Biology 54, 916–935 (2005).

35. B. D. Ortman, A. Bucklin, F. Pages, M. Youngbluth, DNA barcoding the medusozoa using mtCOI. Deep Sea Research Part II: Topical Studies in Oceanography 57, 2148– 2156 (2010).

36. Linné Carl von, Gmelin Johann Friedrich, Bernuset Pierre, Delamollière Jean Baptiste, Caroli a Linné, Systema Naturæ Per Regna Tria Naturæ, Secundum Classes, Ordines, Genera, Species; Cum Characteribus, Differentiis, Synonymis, Locis (1788)vols. I, Part VI.

37. La Martinière, Sur quelques insects. Observations et Mémoires sur la Physique, sur l’Histoire Naturelle et sur les Arts et Métiers tome 31, Pl. II (1787).

38. Péron François, Freycinet Louis Claude Desaulses de, Lesueur Charles Alexandre, Petit Nicolas-Martin, Péron François, Baudin Nicolas, Voyage de Découvertes Aux Terres Australes (Paris, De l’Imprimerie impériale, 1807)vols. Historique, t. 1.

39. C. Chun, Die Siphonophoren der Plankton-Expedition. Ergebnisse der in dem Atlantischen Ocean von Mitte Juli bis Anfang November Bd. 2.K.b, 1–126 (1897).

40. K. C. Schneider, Mittheilungen über Siphonophoren. III. Systematische und andere Bemerkungen. Zoologischer Anzeiger Bd. 21, 51–53, 73–93, 114–133, 153–173, 185–200 (1898).

41. H. Sloane, A Voyage to the Islands Madera, Barbados, Nieves, s. Christophers and Jamaica, with the Natural History of the Herbs and Trees, Four-Footed Beasts, Fishes, Birds, Insects, Reptiles, Etc. Of the Last of Those Islands. (London, 1725)vol. 2.

42. National Oceanographic and Atmospheric Administration. National Weather Service, United States Army. arcGIS user: dhwilso1_NASA., Major ocean currents map.

43. G. E. Hutchinson, The paradox of the plankton. The American Naturalist 95, 137–145 (1961).

44. M. J. Behrenfeld, R. O’Malley, E. Boss, L. Karp-Boss, C. Mundt, Phytoplankton biodiversity and the inverted paradox. ISME Communications 1, 52 (2021).

45. J. Chen, S. Dellaporta, “Urea-based plant DNA miniprep” in The Maize Handbook (Springer, 1994), pp. 526–527.

46. A. Guang, M. Howison, F. Zapata, C. Lawrence, C. W. Dunn, Revising transcriptome assemblies with phylogenetic information. PLOS One 16, e0244202 (2021).

47. S. Koren, B. P. Walenz, K. Berlin, J. R. Miller, N. H. Bergman, A. M. Phillippy, Canu: Scalable and accurate long-read assembly via adaptive k-mer weighting and repeat separation. Genome Research 27, 722–736 (2017).

48. S. Koren, A. Rhie, B. P. Walenz, A. T. Dilthey, D. M. Bickhart, S. B. Kingan, S. Hiendleder, J. L. Williams, T. P. Smith, A. M. Phillippy, De novo assembly of haplotype-resolved genomes with trio binning. Nature Biotechnology 36, 1174–1182 (2018).

49. S. Nurk, B. P. Walenz, A. Rhie, M. R. Vollger, G. A. Logsdon, R. Grothe, K. H. Miga, E. E. Eichler, A. M. Phillippy, S. Koren, HiCanu: Accurate assembly of segmental duplications, satellites, and allelic variants from high-fidelity long reads. Genome Research 30, 1291–1305 (2020).

50. H. Li, Minimap2: Pairwise alignment for nucleotide sequences. Bioinformatics 34, 3094–3100 (2018).

51. M. J. Roach, S. A. Schmidt, A. R. Borneman, Purge Haplotigs: Allelic contig reassignment for third-gen diploid genome assemblies. BMC Bioinformatics 19, 1–10 (2018).

52. L. Coombe, J. X. Li, T. Lo, J. Wong, V. Nikolic, R. L. Warren, I. Birol, LongStitch: High-quality genome assembly correction and scaffolding using long reads. BMC Bioinformatics 22, 1–13 (2021).

53. J. M. Flynn, R. Hubley, C. Goubert, J. Rosen, A. G. Clark, C. Feschotte, A. F. Smit, RepeatModeler2 for automated genomic discovery of transposable element families. Proceedings of the National Academy of Sciences 117, 9451–9457 (2020).

54. M. Manni, M. R. Berkeley, M. Seppey, F. A. Simão, E. M. Zdobnov, BUSCO update: Novel and streamlined workflows along with broader and deeper phylogenetic coverage for scoring of eukaryotic, prokaryotic, and viral genomes. Molecular biology and evolution 38, 4647–4654 (2021).

55. A. M. Bolger, M. Lohse, B. Usadel, Trimmomatic: A flexible trimmer for Illumina sequence data. Bioinformatics 30, 2114–2120 (2014).

56. D. E. Wood, J. Lu, B. Langmead, Improved metagenomic analysis with Kraken 2. Genome Biology 20, 1–13 (2019).

57. C. C. Chang, C. C. Chow, L. C. Tellier, S. Vattikuti, S. M. Purcell, J. J. Lee, Second-generation PLINK: Rising to the challenge of larger and richer datasets. Gigascience 4, s13742–015 (2015).

58. H. Li, Aligning sequence reads, clone sequences and assembly contigs with BWA-MEM. arXiv (2013).

59. H. Li, A statistical framework for SNP calling, mutation discovery, association mapping and population genetical parameter estimation from sequencing data. Bioinformatics 27, 2987–2993 (2011).

60. T. R. Ranallo-Benavidez, K. S. Jaron, M. C. Schatz, GenomeScope 2.0 and Smudgeplot for reference-free profiling of polyploid genomes. Nature Communications 11, 1432 (2020).

61. G. Marcais, C. Kingsford, Jellyfish: A fast k-mer counter. Tutorialis e Manuais 1, 1038 (2012).

62. J.-J. Jin, W.-B. Yu, J.-B. Yang, Y. Song, C. W. DePamphilis, T.-S. Yi, D.-Z. Li, GetOrganelle: A fast and versatile toolkit for accurate de novo assembly of organelle genomes. Genome Biology 21, 1–31 (2020).

63. K. Katoh, D. M. Standley, MAFFT multiple sequence alignment software version 7: Improvements in performance and usability. Molecular biology and evolution 30, 772– 780 (2013).

64. B. Minh, H. Schmidt, O. Chernomor, D. Schrempf, M. Woodhams, A. Von Haeseler, R. L. IQ-TREE, IQ-TREE2: New models and efficient methods for phylogenetic inference in the genomic era. DOI:10.1093/molbev/msaa015 37, 1530–1534 (2020).

65. S. Kalyaanamoorthy, B. Q. Minh, T. K. Wong, A. Von Haeseler, L. S. Jermiin, ModelFinder: Fast model selection for accurate phylogenetic estimates. Nature Methods 14, 587–589 (2017).

66. D. T. Hoang, O. Chernomor, A. Von Haeseler, B. Q. Minh, L. S. Vinh, UFBoot2: Improving the ultrafast bootstrap approximation. Molecular Biology and Evolution 35, 518–522 (2018).

67. J. Chifman, L. Kubatko, Quartet inference from SNP data under the coalescent model. Bioinformatics 30, 3317–3324 (2014).

68. T. S. Korneliussen, A. Albrechtsen, R. Nielsen, ANGSD: Analysis of Next Generation Sequencing Data. BMC Bioinformatics 15, 356 (2014).

69. J. Meisner, A. Albrechtsen, Inferring population structure and admixture proportions in low-depth NGS data. Genetics 210, 719–731 (2018).

70. K. L. Korunes, K. Samuk, Pixy: Unbiased estimation of nucleotide diversity and divergence in the presence of missing data. Molecular Ecology Resources 21, 1359–1368 (2021).

71. M. Malinsky, M. Matschiner, H. Svardal, Dsuite-fast D-statistics and related admixture evidence from VCF files. Molecular Ecology Resources 21, 584–595 (2021).

