## Supplementary material for "Global genomics of the man-o’-war (*Physalia*) reveals biodiversity at the ocean surface": supp. figures and tables

### Supplementary tables and figures

**4 Phylogeny inferred from SNP data** **8**

**5 Test of introgression** **9**

**6 Individual genetree** **10**

**7 Morphological classification** **14**

**8 Subpopulation analyses** **15**

### 1 Genome assembly statistics

**Table S1** : Genome assembly statistics.

| measurement | primary | alternate |
|---|---|---|
| total scaffolds | 2,386 | 5,480 |
| total contigs | 3,646 | 9,440 |
| scaffold total size, GB | 3.33 | 2.69 |
| scaffold N50, MB | 10.4 | 4.6 |
| scaffolds >10MB | 96 | 31 |
| scaffolds >1MB | 449 | 625 |

**Table S2** : Genome BUSCO statistics.

| measurement | primary | alternate |
|---|---|---|
| BUSCO complete | 89.7% | 86.9% |
| single-copy | 84.8% | 80.5% |
| duplicated | 4.9% | 6.4% |
| fragmented | 5.3% | 5.6% |
| missing | 5.0% | 7.5% |

#### 1.1 Genome size estimate

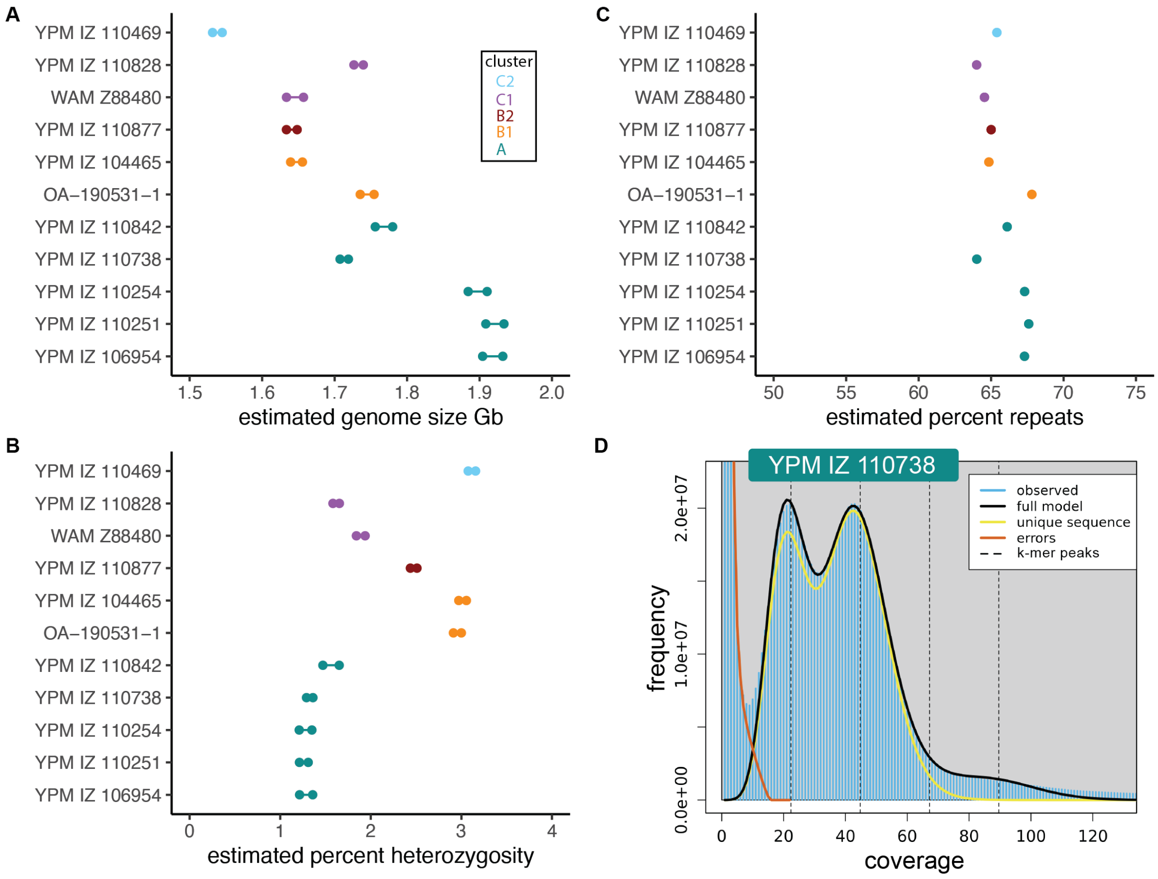

**Figure S1** : Genome statistics for samples sequenced to a depth sufficient for a good model fit with a k-mer approach. A, genome size estimates from `GenomeScope`, in gigabases. Colors indicate clusters as in Fig. 2. B, estimated genome percent heterozygosity. C, estimated percent of the genome that is repeat sequences. D, example `GenomeScope model` fit for one Atlantic specimen.

#### 2 Population genomics

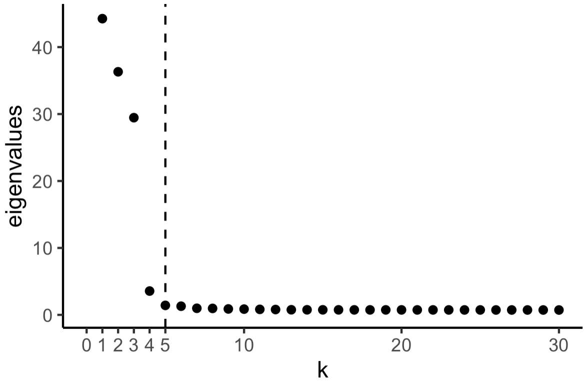

**Figure S2** : Eigenvalues of the covariance matrix. The optimal number of components (k=5), as determined with PCANGSD, is shown with a dotted line.

##### 2.1 Expanded sample set

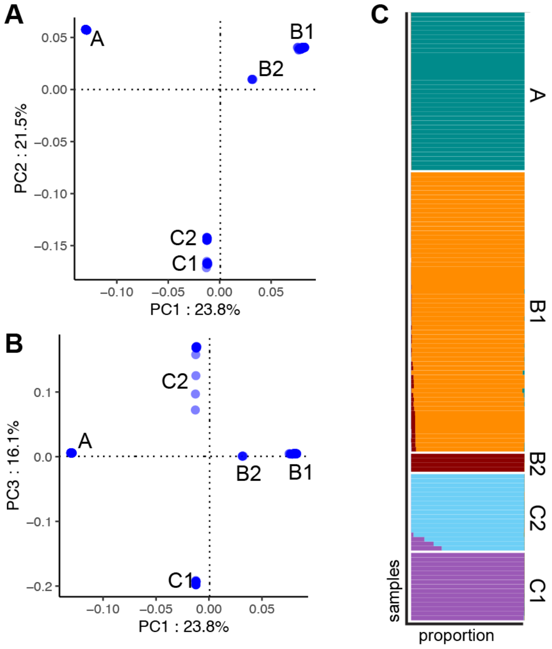

**Figure S3** : Principal component analysis (PCA, A-B) and shared ancestry analysis (C) of 133 high and moderate quality samples, mapping reads to non-repeat regions of the reference genome.

#### 2.2 Iso-Seq reference

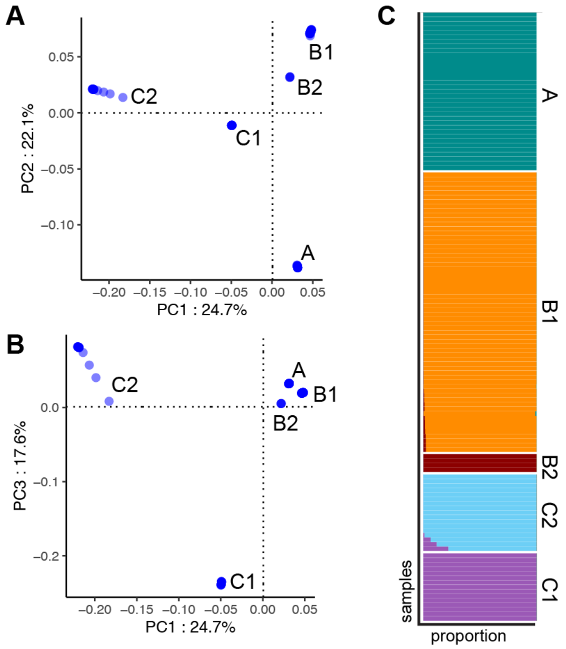

**Figure S4** : PCA (A-B) and shared ancestry analysis (C) of 133 samples, mapping reads to a reference transcriptome assembled from Iso-Seq data.

### 3 Population statistics

#### 3.1 Estimates of *Fst*

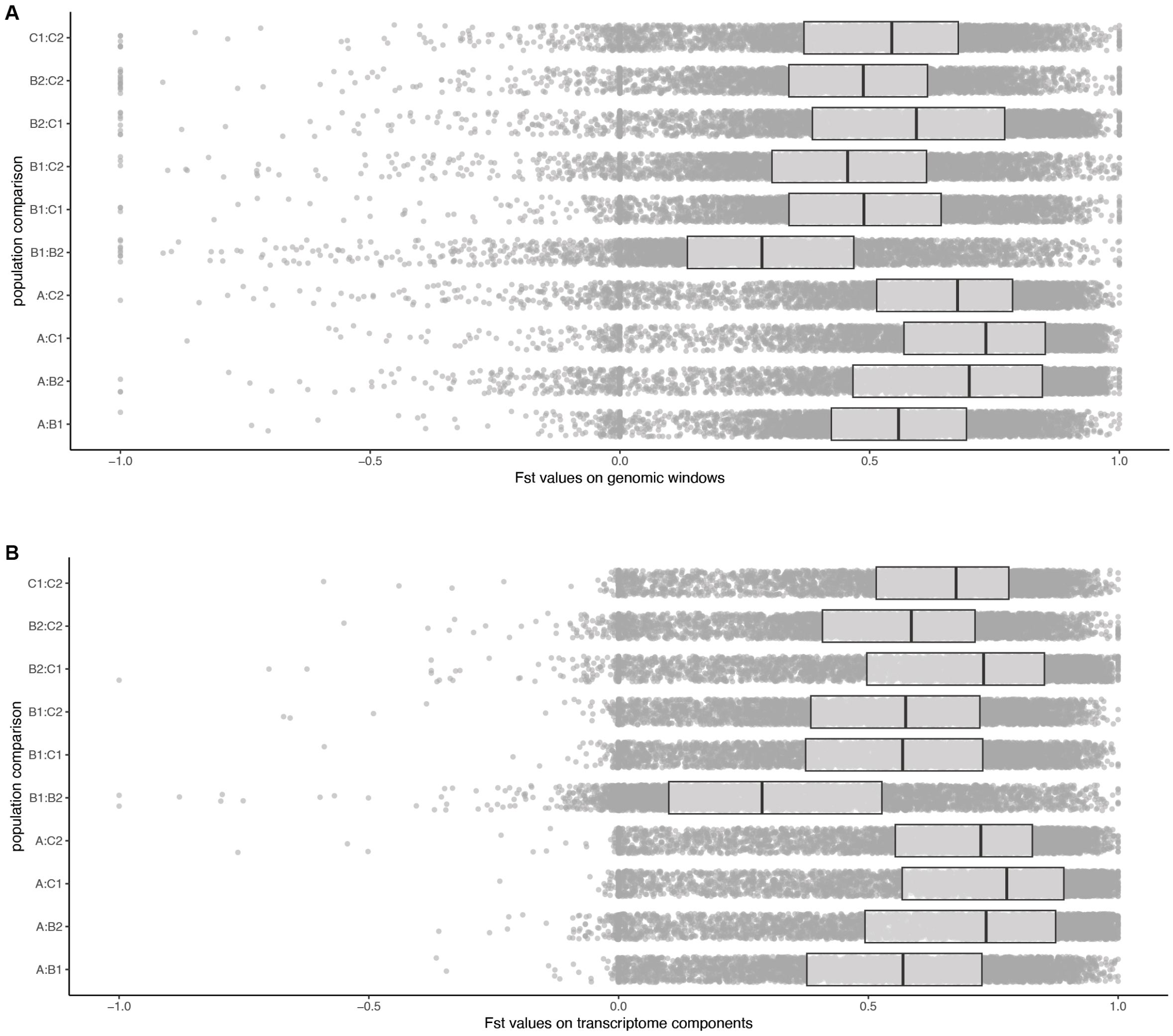

**Figure S5** : Genomic differentiation index *Fst* values between clusters as defined in Figs. 1-2, box-plots show mean and quartile values. A, *Fst* values across 5,000 randomly sampled non-repeat genomic windows. B, *Fst* values across 5,000 randomly sampled transcripts.

#### 3.2 Estimates of *pi*

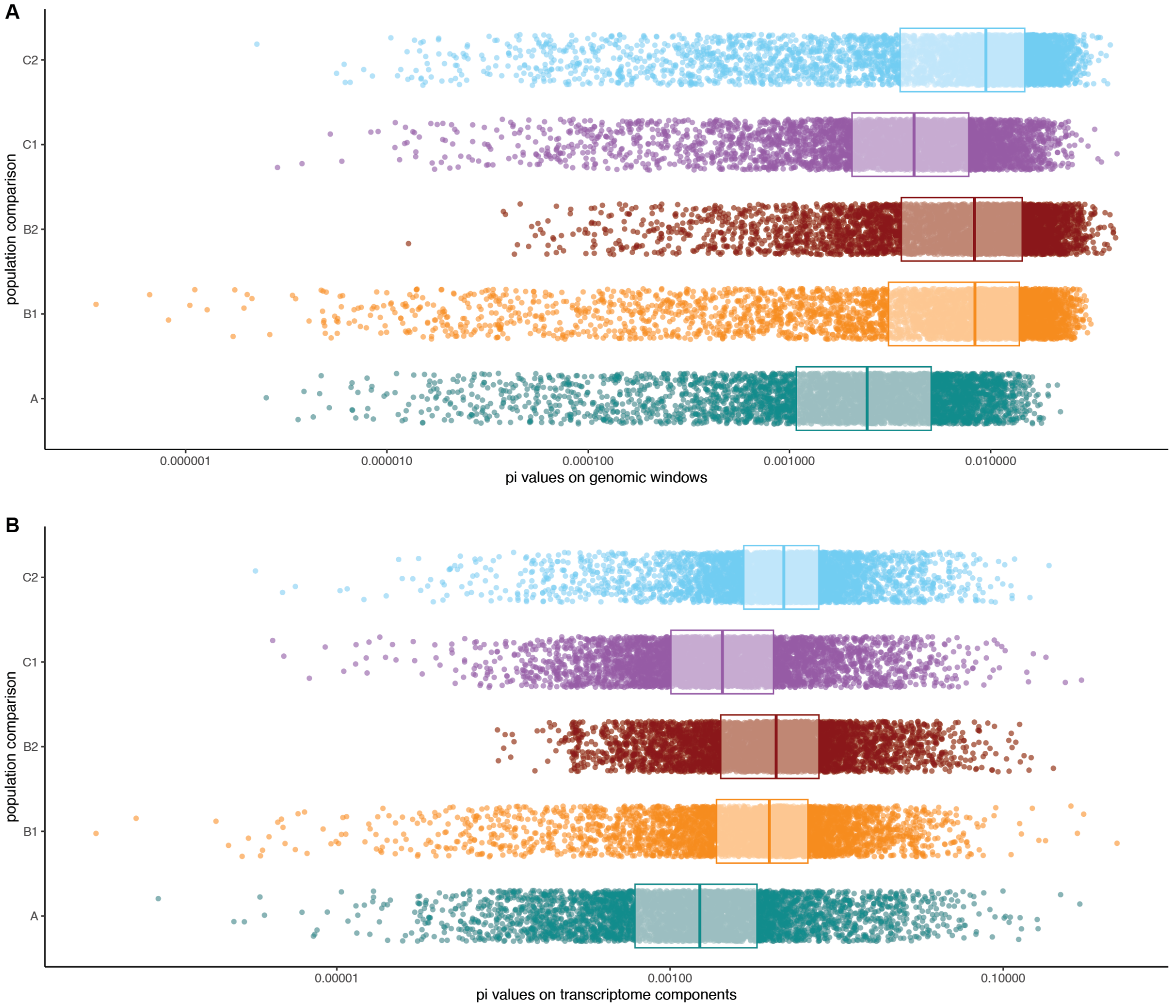

**Figure S6** : Nucleotide diversity index *pi* values between clusters, as defined in Figs. 1-2, box-plots show mean and quartile values. A, *pi* values across 5,000 randomly sampled non-repeat genomic windows. B, *pi* values across 5,000 randomly sampled transcripts.

### 4 Phylogeny inferred from SNP data

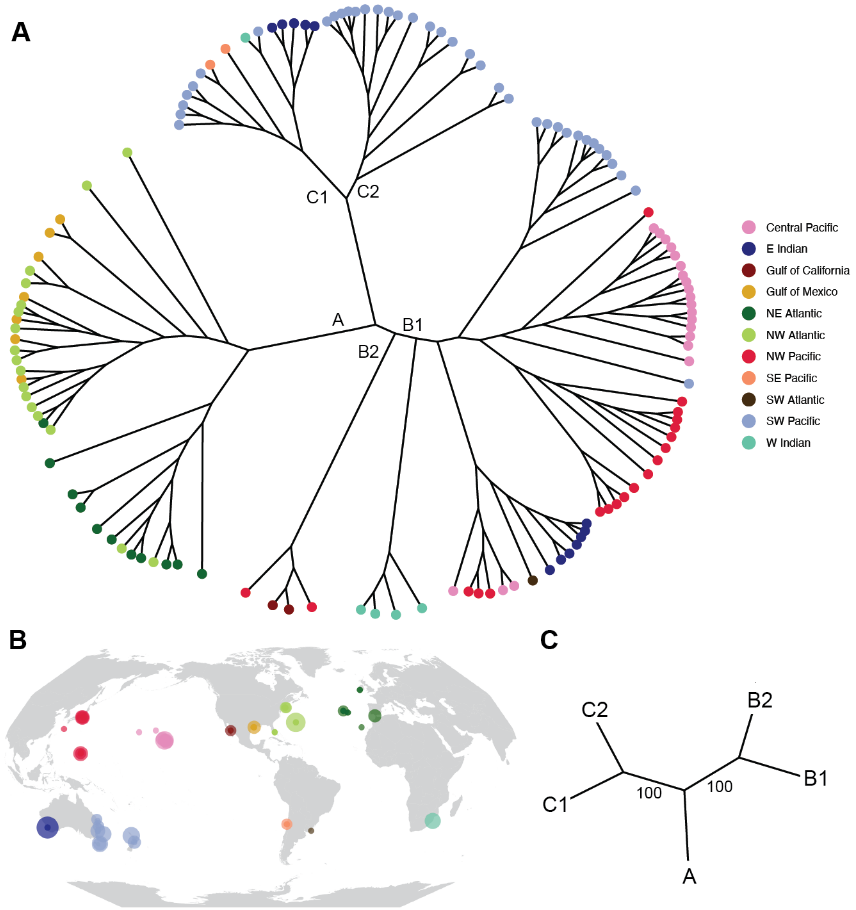

**Figure S7** : A, unrooted phylogeny of specimens, inferred with `SVDQuartets` on ~800k high-quality SNPs from non-repeat genomic regions. Internal branches are annotated with cluster labels, as defined in Figs. 1-2. Colors indicate region, as in (B). C, unrooted species phylogeny using the same SNP data. Coalescent bootstrap values are shown at internal branches.

### 5 Test of introgression

**Table S3** : D-statistics between quartets of lineages

| P1 | P2 | P3 | P4 | D-statistic | Z score | p-value |
|---|---|---|---|---|---|---|
| B2 | B1 | A | C1 | 0.018 | 1.665 | 0.096 |
| B2 | B1 | A | C2 | 0.005 | 0.489 | 0.625 |
| B1 | A | C1 | C2 | 0.012 | 0.902 | 0.367 |
| B2 | A | C1 | C2 | 0.003 | 0.302 | 0.762 |
| B1 | B2 | C1 | C2 | 0.015 | 1.377 | 0.169 |

### 6 Individual genetree

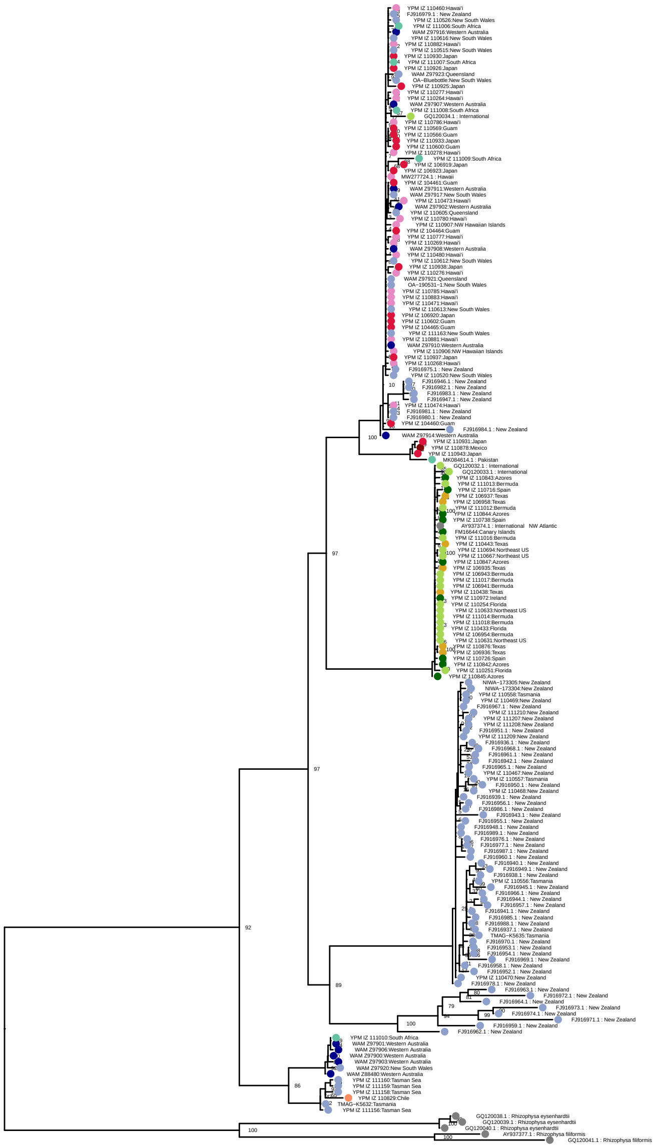

**Figure S8** : Phylogeny of cytochrome oxidase 1 (CO1) sequences, assembled using *in silico* PCR from reads, and publicly available sequences from NCBI. Colors indicate region, as in Fig. S7B. Bootstrap support values shown at nodes. Rooted using *Rhizophysa* as an outgroup.

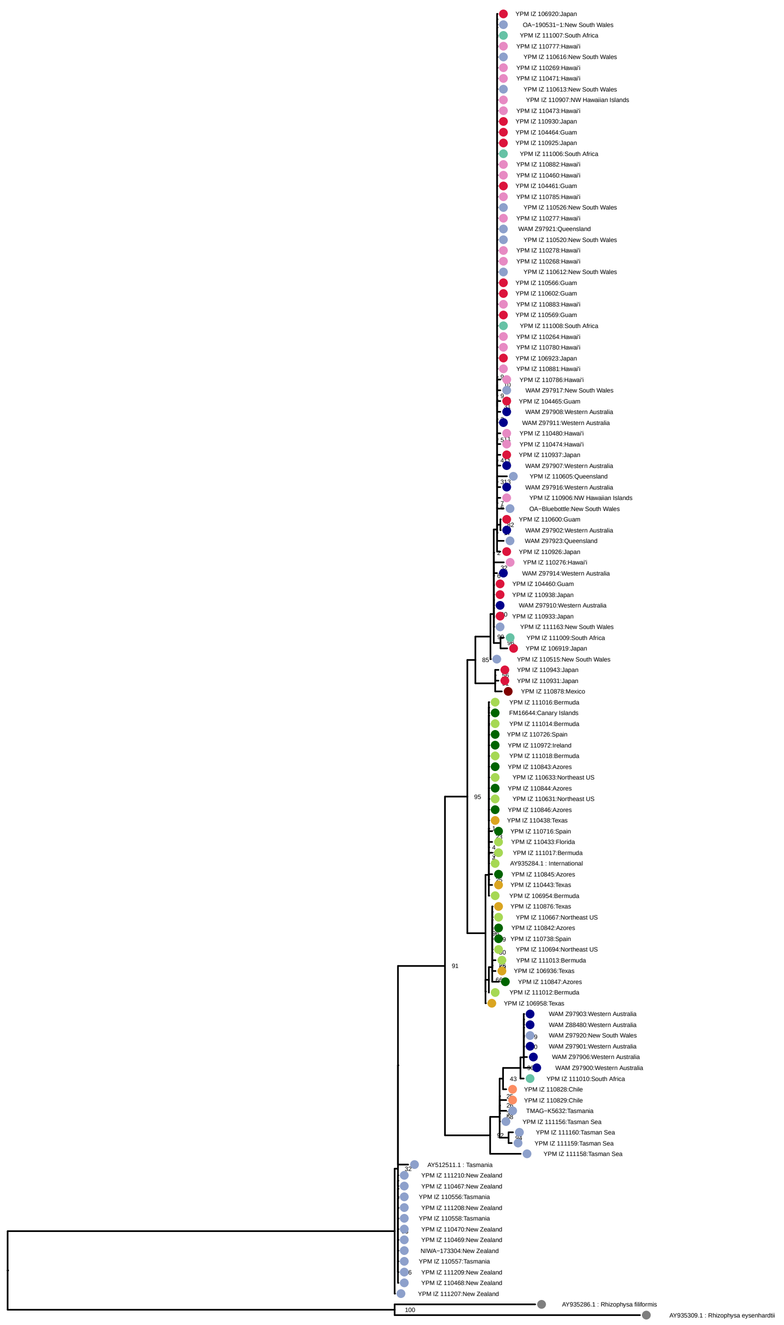

**Figure S9** : Phylogeny of 16S ribosomal RNA sequences, assembled using *in silico* PCR from reads, and publicly available sequences from NCBI. Colors indicate region, as in Fig. S7B. Bootstrap support values shown at nodes. Rooted using *Rhizophysa* as an outgroup.

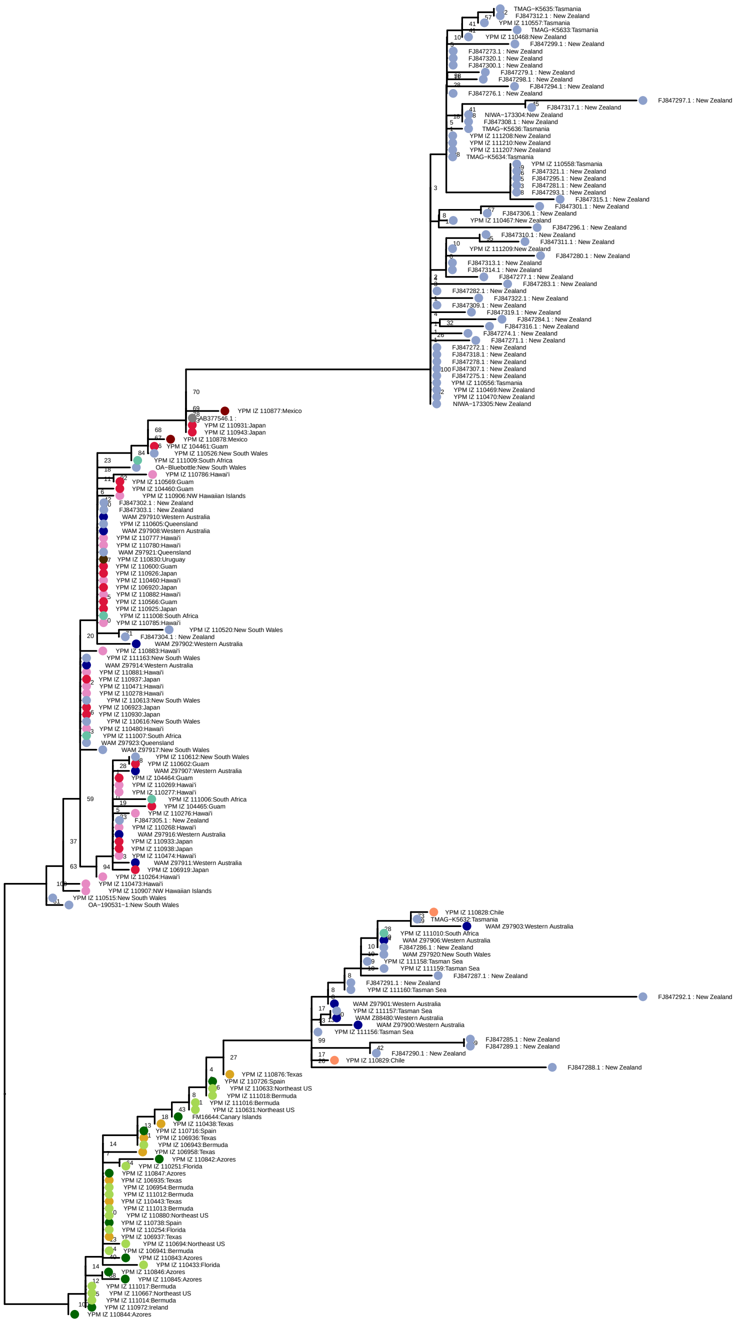

**Figure S10** : Unrooted phylogeny of Internal Transcribed Spacer (ITS) sequences, assembled using *in silico* PCR from reads, and publicly available sequences from NCBI. Colors indicate region, as in Fig. S7B. Bootstrap support values shown at nodes.

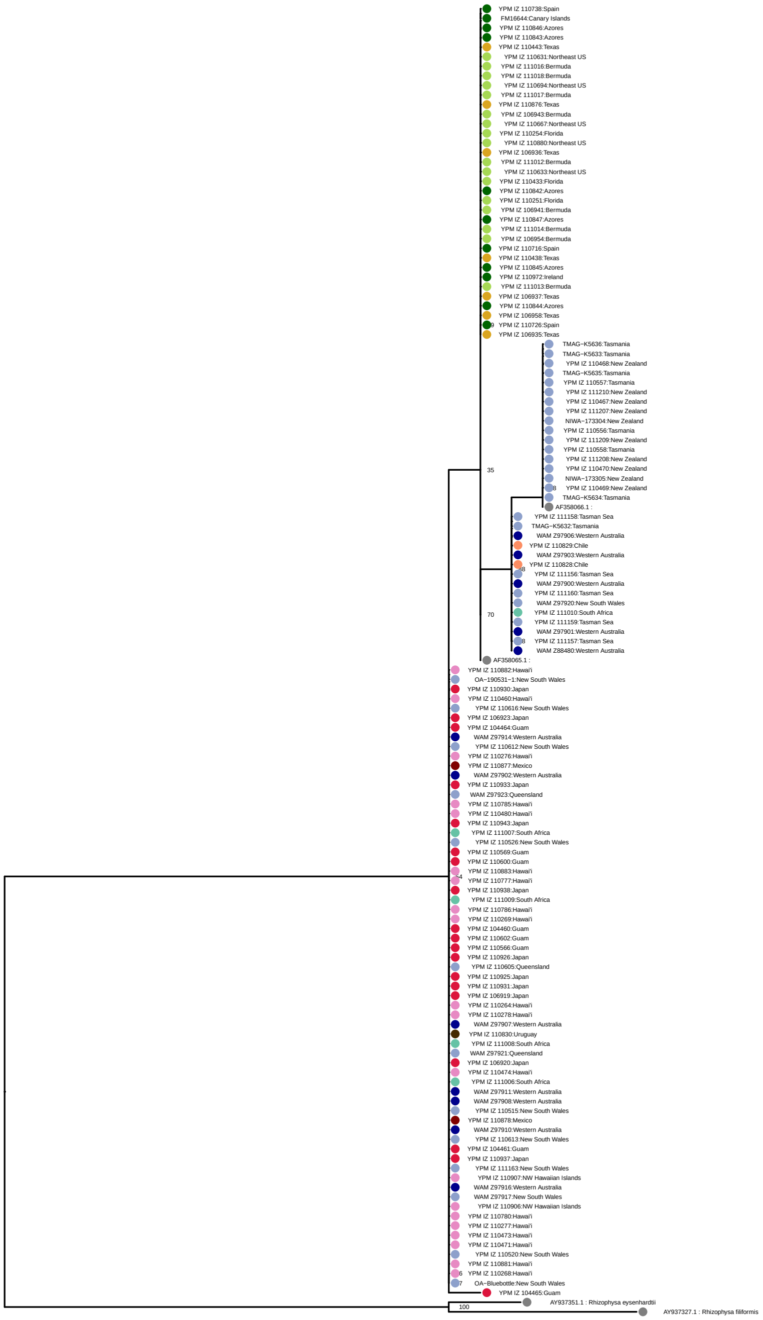

**Figure S11** : Phylogeny of 18S ribosomal RNA sequences, assembled using *in silico* PCR from reads, and publicly available sequences from NCBI. Colors indicate region, as in Fig. S7B. Bootstrap support values shown at nodes. Rooted using *Rhizophysa* as an outgroup.

### 7 Morphological classification

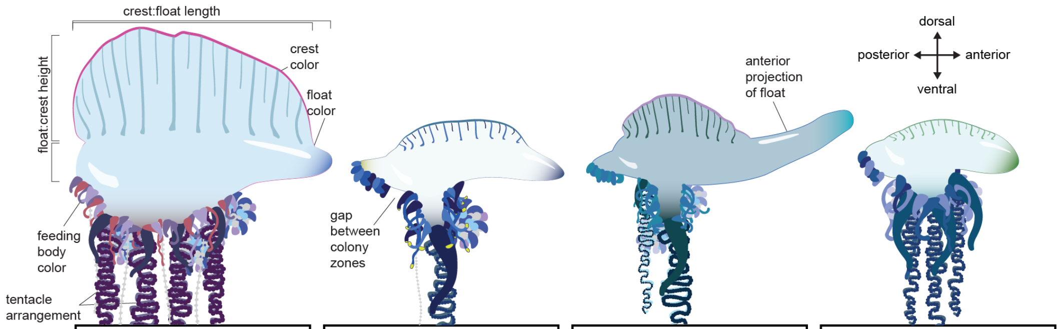

| | *P. physalis* | | *P. utriculus* | | *P. megalista* | | *P.* sp C2 | |
|---|---|---|---|---|---|---|---|---|
| feeding body color | reddish | yellow-tipped | yellow-tipped | reddish | | reddish OR yellow-tipped | | |
| float color | | clear, glassy | clear, glassy | | | clear, glassy | | |
| crest color | | | | | purple | | | pink, purple |
| crest:float height | as tall as float | | | | | | no visible height | |
| float:crest length | anterior projection <1/4 crest length | anterior projection >3/4 crest length | | anterior projection >3/4 crest length | anterior projection >3/4 crest length | anterior projection <1/4 crest length | | |
| gap between growth zones | | present | present | | present | | | |
| arrangement principle tentacles | >2 central | | 1-2 central | >2 central | | >2 central | >2 central | |

strong indicator, sufficient to confirm

weak indicator, must all be present to confirm

counterindicator, sufficient to exclude

**Figure S12** : Rules-based analysis for positive identification of iNaturalist images based on scored traits. Images of poor quality or of specimens scored as having juvenile characteristics (e.g., globular float, few zooids) were excluded. Rules were selected to classify high-confidence observations of adults from each morphology, and to minimize overlap between them. Individual specimens of each morphology may deviate from these characters (e.g., if the sail is not raised at the time of observation).

### 8 Subpopulation analyses

#### 8.1 Cluster B1+B2

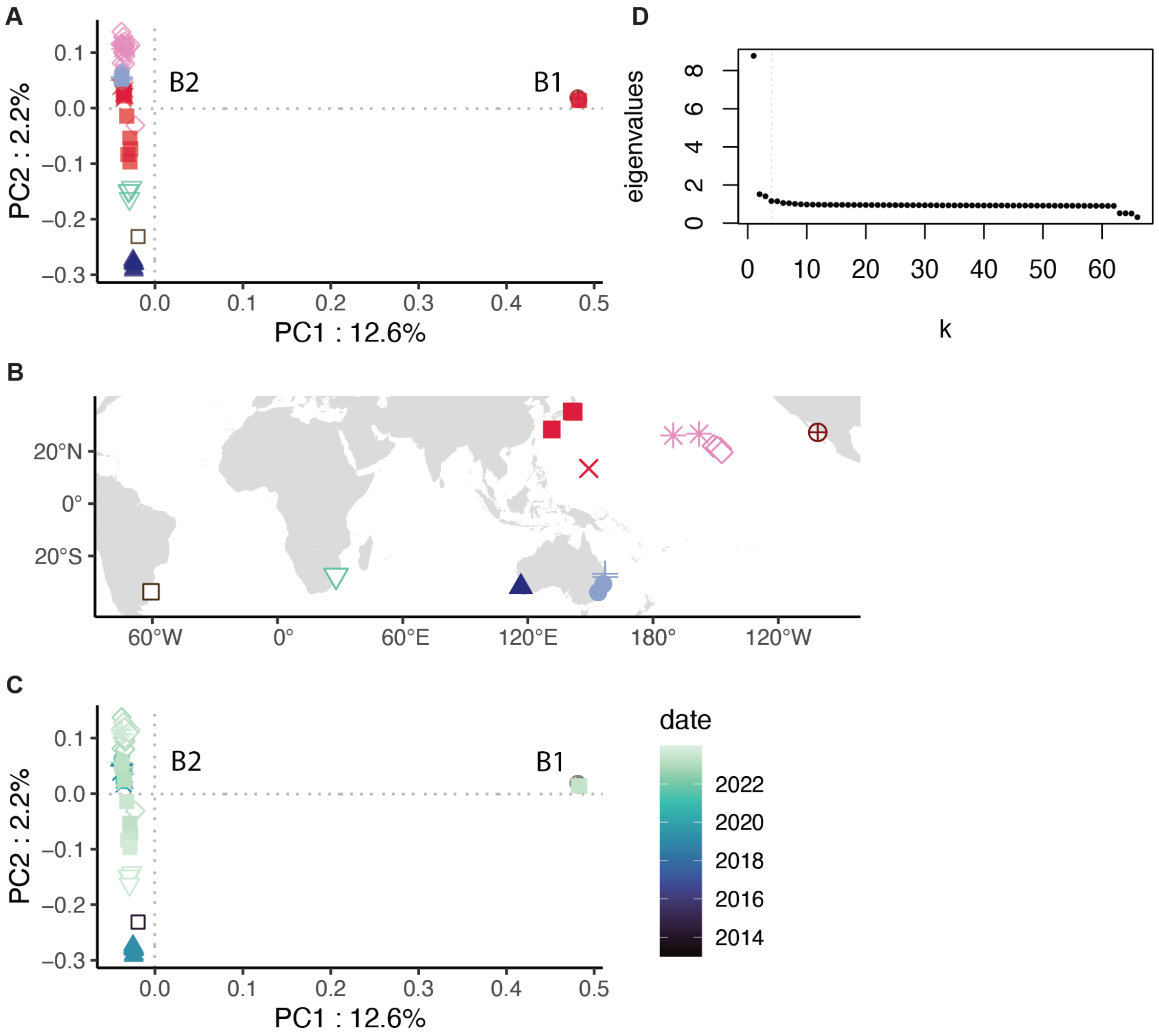

**Figure S13** : A, PCA of samples in clusters B1 and B2 together. B, map of samples. C, PCA, colored by collection date. D, eigenvalues of the covariance matrix.

#### 8.2 Temporal variation

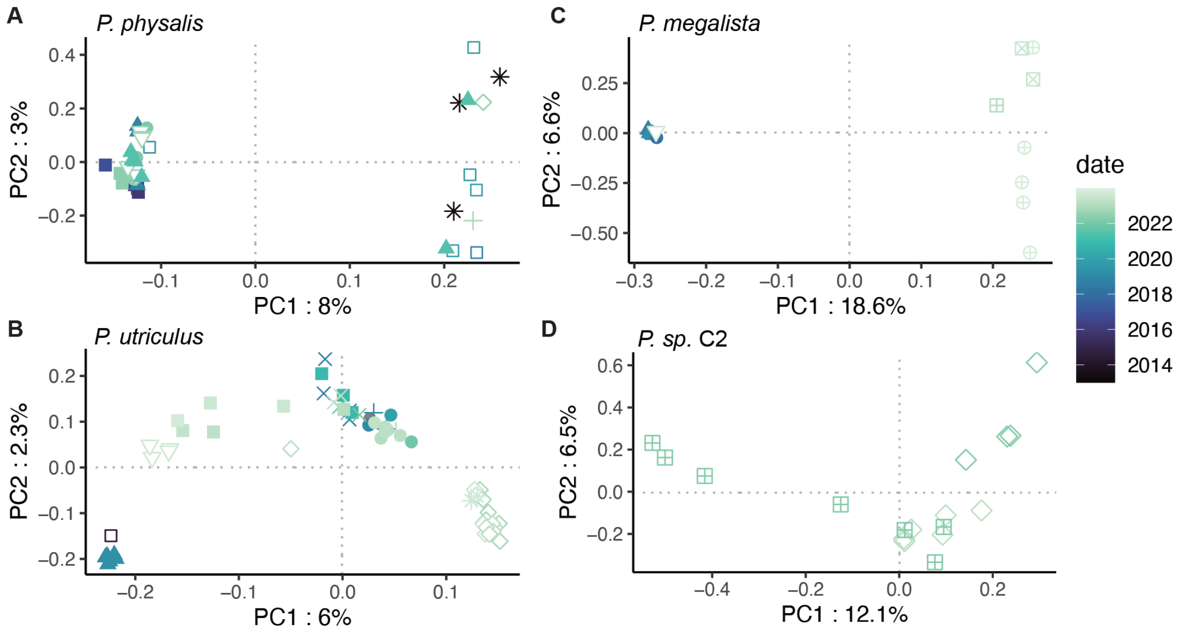

**Figure S14** : A-D, Sample collection date, visualized on the first two PCs of genomic variation for each species. Shapes correspond to geographic region, as in Fig 4.

#### 8.3 K-means clustering subclustering

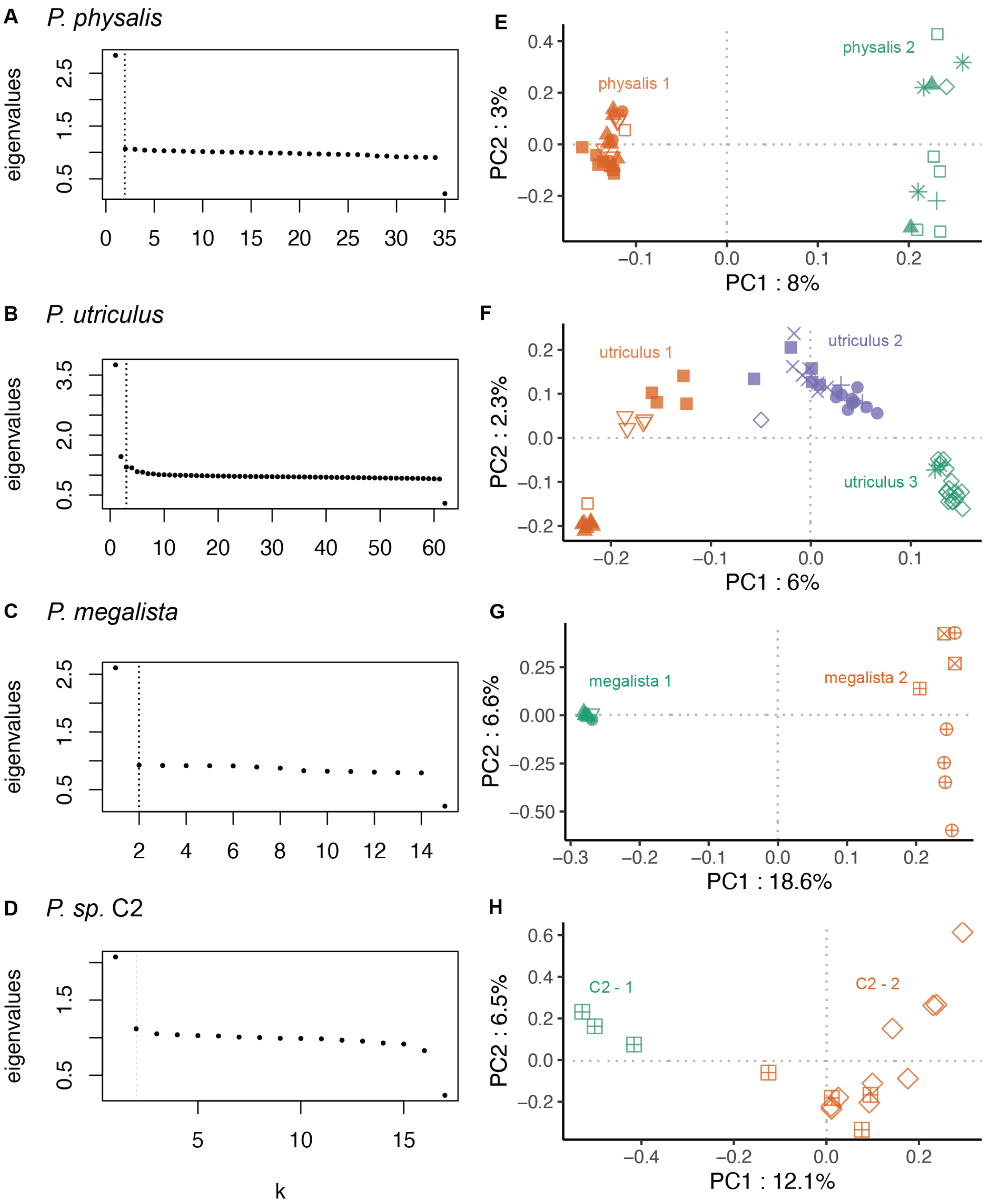

**Figure S15** : A-D, eigenvalues of the covariance matrix, the optimal number of clusters marked by a dashed line. E-H,Result of k-means clustering within each of the four species. Shapes correspond to geographic region, as in Fig. 4

#### 8.4 *Fst*, *pi*, and *Dxy* values in subpopulations

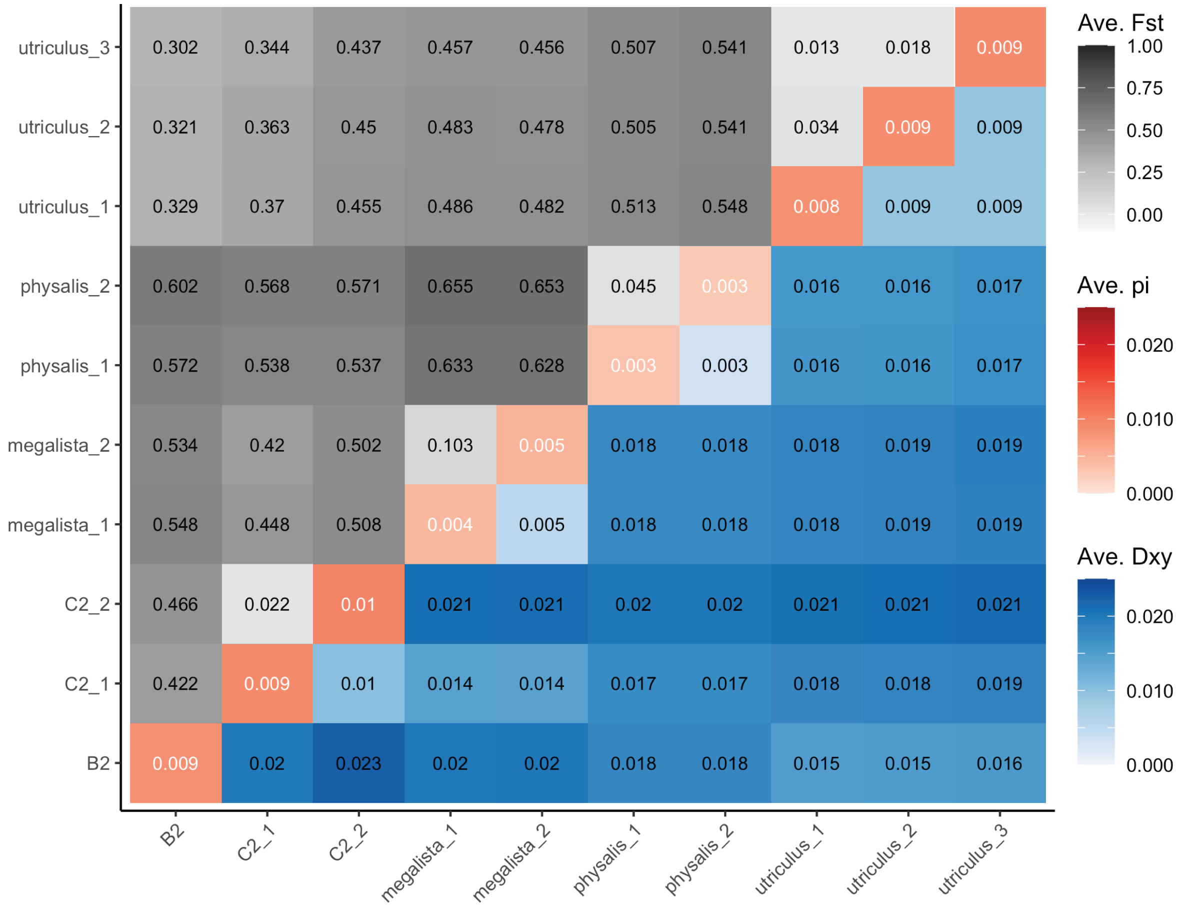

**Figure S16** : Reciprocal average *Fst* (gray), *pi* (red), and *Dxy* (blue) values between subpopulations, defined by k-means clustering of the covariance matrix within populations. Lineage B2 was treated as one population.

#### 8.5 *Fst*, *pi*, and *Dxy* values in lineages and regions

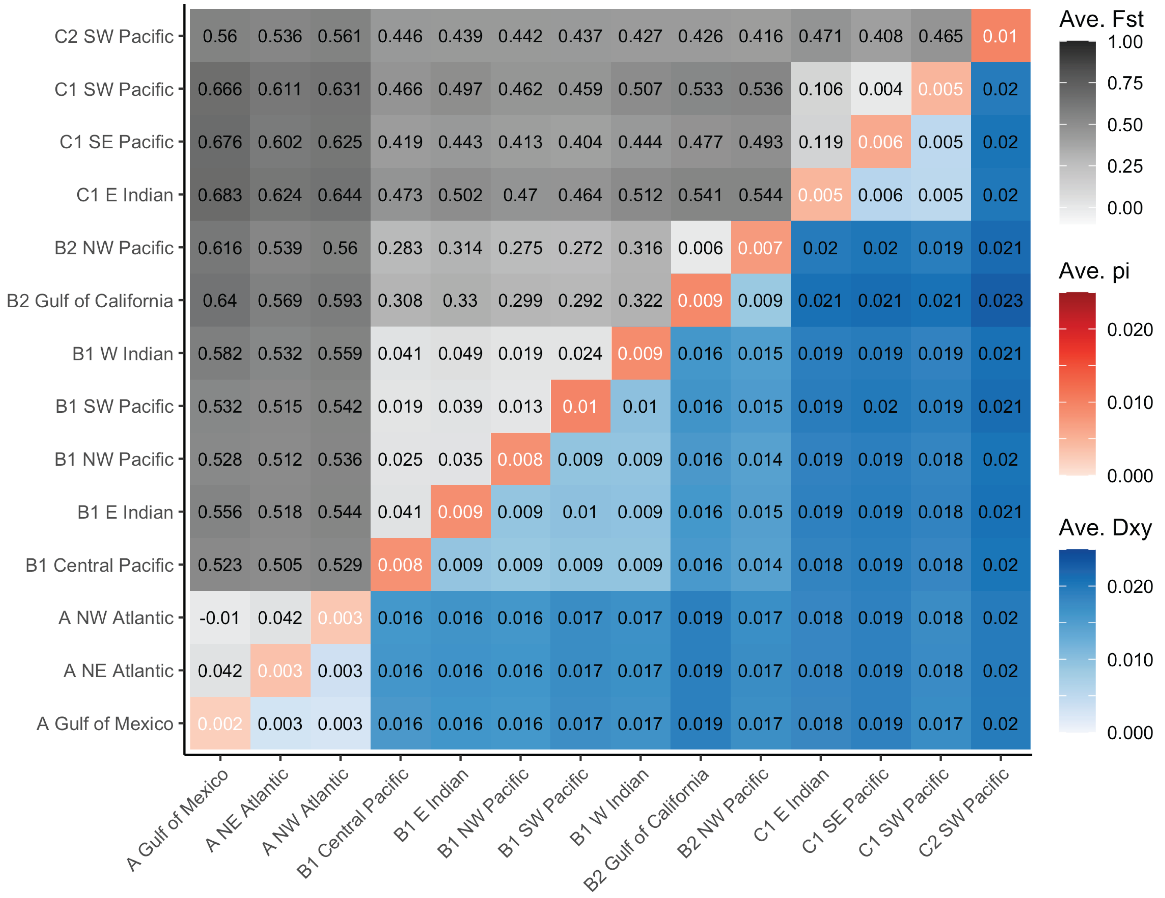

**Figure S17** : Reciprocal average *Fst* (gray), *pi* (red), and *Dxy* (blue) values between subpopulations, defined by lineage and oceanic region, and excluding groupings with only one representative sample.
