## Supplementary material for "Global genomics of the man-o’-war (*Physalia*) reveals biodiversity at the ocean surface": supp. specimen information

### Supplementary material: specimen collection information

This document contains collection information and photos for samples intended for population genomic sequencing.

Table 1: Specimen data.

| ID | cluster | ocean region | location | latitude | longitude | date collected | quality status | permit |
|---|---|---|---|---|---|---|---|---|
| FM16644 | A | NE Atlantic | Canary Islands | 28.591940 | -17.920910 | 5/14/2023 | high quality | |
| TMAG K5632 | C1 | SW Pacific | Tasmania | -41.863900 | 148.286500 | 1/14/2023 | moderate quality | |
| TMAG K5633 | C2 | SW Pacific | Tasmania | -41.863900 | 148.286500 | 1/14/2023 | moderate quality | |
| TMAG K5634 | C2 | SW Pacific | Tasmania | -41.863900 | 148.286500 | 1/14/2023 | moderate quality | |
| TMAG K5635 | C2 | SW Pacific | Tasmania | -41.863900 | 148.286500 | 1/14/2023 | moderate quality | |
| TMAG K5636 | C2 | SW Pacific | Tasmania | -41.863900 | 148.286500 | 1/14/2023 | moderate quality | |
| OA 190531 1 | B1 | SW Pacific | New South Wales | -33.891500 | 151.276700 | 12/15/2019 | high quality | |
| OA Bluebottle | B1 | SW Pacific | New South Wales | -33.920300 | 151.258100 | 1/1/2022 | moderate quality | |
| WAMZ88480 | C1 | E Indian | Western Australia | -31.908144 | 115.754811 | 8/3/2018 | high quality | |
| WAMZ97900 | C1 | E Indian | Western Australia | -31.855786 | 115.751741 | 4/19/2019 | high quality | |
| WAMZ97901 | C1 | E Indian | Western Australia | -31.855786 | 115.751741 | 4/19/2019 | high quality | |
| WAMZ97902 | B1 | E Indian | Western Australia | -31.855786 | 115.751741 | 4/19/2019 | high quality | |
| WAMZ97903 | C1 | E Indian | Western Australia | -31.855786 | 115.751741 | 4/19/2019 | high quality | |
| WAMZ97906 | C1 | E Indian | Western Australia | -31.855786 | 115.751741 | 4/19/2019 | high quality | |
| WAMZ97907 | B1 | E Indian | Western Australia | -31.855786 | 115.751741 | 4/19/2019 | high quality | |
| WAMZ97908 | B1 | E Indian | Western Australia | -31.855786 | 115.751741 | 4/19/2019 | high quality | |
| WAMZ97910 | B1 | E Indian | Western Australia | -31.855786 | 115.751741 | 4/19/2019 | high quality | |
| WAMZ97911 | B1 | E Indian | Western Australia | -31.855786 | 115.751741 | 4/19/2019 | high quality | |
| WAMZ97914 | B1 | E Indian | Western Australia | -31.855786 | 115.751741 | 4/19/2019 | high quality | |
| WAMZ97916 | B1 | E Indian | Western Australia | -31.855786 | 115.751741 | 4/19/2019 | high quality | |
| WAMZ97917 | B1 | SW Pacific | New South Wales | -33.787002 | 151.288609 | 1/18/2019 | high quality | |
| WAMZ97920 | C1 | SW Pacific | New South Wales | -33.987424 | 151.231329 | 2/5/2018 | high quality | |
| WAMZ97921 | B1 | SW Pacific | Queensland | -26.784003 | 153.140596 | 1/19/2019 | high quality | |
| WAMZ97923 | B1 | SW Pacific | Queensland | -26.784003 | 153.140596 | 1/19/2019 | high quality | |
| YPM IZ 104460 | B1 | NW Pacific | Guam | 13.427861 | 144.798518 | 11/23/2018 | high quality | Guam Department of Agriculture, Division of Aquatic and Wildlife Resources |
| YPM IZ 104461 | B1 | NW Pacific | Guam | 13.427861 | 144.798518 | 11/23/2018 | high quality | Guam Department of Agriculture, Division of Aquatic and Wildlife Resources |
| YPM IZ 104464 | B1 | NW Pacific | Guam | 13.427861 | 144.798518 | 11/23/2018 | high quality | Guam Department of Agriculture, Division of Aquatic and Wildlife Resources |
| YPM IZ 104465 | B1 | NW Pacific | Guam | 13.427861 | 144.798518 | 11/23/2018 | high quality | Guam Department of Agriculture, Division of Aquatic and Wildlife Resources |
| YPM IZ 106919 | B1 | NW Pacific | Japan | 35.307390 | 139.480651 | 11/19/2020 | high quality | |
| YPM IZ 106920 | B1 | NW Pacific | Japan | 35.307390 | 139.480651 | 11/19/2020 | high quality | |
| YPM IZ 106923 | B1 | NW Pacific | Japan | 35.307390 | 139.480651 | 11/19/2020 | high quality | |
| YPM IZ 106935 | A | Gulf of Mexico | Texas | 29.19532 | -94.8429383 | 3/2/2016 | high quality | |
| YPM IZ 106936 | A | Gulf of Mexico | Texas | 29.19532 | -94.8429383 | 3/3/2016 | high quality | |
| YPM IZ 106937 | A | Gulf of Mexico | Texas | 29.19532 | -94.8429383 | 3/3/2016 | high quality | |
| YPM IZ 106941 | A | NW Atlantic | Bermuda | 32.323333 | -64.656300 | 5/18/2019 | high quality | Bermuda Department of Environment and Natural Resources; SP190501 |
| YPM IZ 106943 | A | NW Atlantic | Bermuda | 32.341117 | -64.669133 | 5/18/2019 | high quality | Bermuda Department of Environment and Natural Resources; SP190501 |
| YPM IZ 106954 | A | NW Atlantic | Bermuda | 32.367233 | -64.714550 | 5/20/2019 | high quality | Bermuda Department of Environment and Natural Resources; SP190501 |
| YPM IZ 106958 | A | Gulf of Mexico | Texas | 29.195320 | -94.842938 | 3/18/2017 | high quality | |
| YPM IZ 110251 | A | NW Atlantic | Florida | 25.737060 | -80.152790 | 12/22/2022 | high quality | |

Table 1: Specimen data. *(continued)*

| ID | cluster | ocean region | location | latitude | longitude | date collected | quality status | permit |
|---|---|---|---|---|---|---|---|---|
| YPM IZ 110254 | A | NW Atlantic | Florida | 25.736470 | -80.153500 | 12/22/2022 | high quality | |
| YPM IZ 110264 | B1 | Central Pacific | Hawai'i | 22.069018 | -159.316685 | 10/31/2022 | high quality | |
| YPM IZ 110268 | B1 | Central Pacific | Hawai'i | 22.069018 | -159.316685 | 11/4/2022 | moderate quality | |
| YPM IZ 110269 | B1 | Central Pacific | Hawai'i | 22.069018 | -159.316685 | 11/4/2022 | high quality | |
| YPM IZ 110276 | B1 | Central Pacific | Hawai'i | 20.9225222 | -156.4926917 | 11/19/2022 | high quality | |
| YPM IZ 110277 | B1 | Central Pacific | Hawai'i | 20.9225222 | -156.4926917 | 11/19/2022 | moderate quality | |
| YPM IZ 110278 | B1 | Central Pacific | Hawai'i | 20.9225222 | -156.4926917 | 11/19/2022 | high quality | |
| YPM IZ 110433 | A | NW Atlantic | Florida | 25.741630 | -80.175194 | 3/14/2023 | high quality | |
| YPM IZ 110438 | A | Gulf of Mexico | Texas | 28.989194 | -95.237778 | 3/24/2023 | moderate quality | Texas NR Saltwater Fishing License, 958064721164 |
| YPM IZ 110443 | A | Gulf of Mexico | Texas | 28.951778 | -95.283167 | 3/24/2023 | high quality | Texas NR Saltwater Fishing License, 958064721164 |
| YPM IZ 110460 | B1 | Central Pacific | Hawai'i | 21.400790 | -157.734500 | 11/14/2022 | high quality | |
| YPM IZ 110467 | C2 | SW Pacific | New Zealand | -41.330278 | 174.831944 | 2/16/2023 | high quality | |
| YPM IZ 110468 | C2 | SW Pacific | New Zealand | -41.330000 | 174.833056 | 2/16/2023 | high quality | |
| YPM IZ 110469 | C2 | SW Pacific | New Zealand | -41.330000 | 174.833056 | 2/16/2023 | high quality | |
| YPM IZ 110470 | C2 | SW Pacific | New Zealand | -41.330000 | 174.833056 | 2/16/2023 | high quality | |
| YPM IZ 110471 | B1 | Central Pacific | Hawai'i | 21.400790 | -157.734500 | 4/9/2023 | high quality | |
| YPM IZ 110473 | B1 | Central Pacific | Hawai'i | 21.400790 | -157.734500 | 4/9/2023 | high quality | |
| YPM IZ 110474 | B1 | Central Pacific | Hawai'i | 21.454600 | -157.748500 | 4/4/2023 | high quality | |
| YPM IZ 110480 | B1 | Central Pacific | Hawai'i | 21.454600 | -157.748500 | 4/6/2023 | high quality | |
| YPM IZ 110515 | B1 | SW Pacific | New South Wales | -33.947500 | 151.257361 | 2/9/2023 | high quality | |
| YPM IZ 110520 | B1 | SW Pacific | New South Wales | -33.947500 | 151.257361 | 2/19/2023 | high quality | |
| YPM IZ 110526 | B1 | SW Pacific | New South Wales | -33.947500 | 151.257361 | 4/23/2023 | high quality | |
| YPM IZ 110556 | C2 | SW Pacific | Tasmania | -41.996839 | 148.284729 | 12/25/2022 | high quality | |
| YPM IZ 110557 | C2 | SW Pacific | Tasmania | -41.996839 | 148.284729 | 12/25/2022 | high quality | |
| YPM IZ 110558 | C2 | SW Pacific | Tasmania | -41.996839 | 148.284729 | 12/25/2022 | high quality | |
| YPM IZ 110566 | B1 | NW Pacific | Guam | 13.605463 | 144.907196 | 1/24/2022 | high quality | Guam Department of Agriculture, Division of Aquatic and Wildlife Resources |
| YPM IZ 110569 | B1 | NW Pacific | Guam | 13.427664 | 144.801336 | 1/27/2022 | moderate quality | Guam Department of Agriculture, Division of Aquatic and Wildlife Resources |
| YPM IZ 110600 | B1 | NW Pacific | Guam | 13.372435 | 144.773357 | 4/27/2023 | high quality | Guam Department of Agriculture, Division of Aquatic and Wildlife Resources |
| YPM IZ 110602 | B1 | NW Pacific | Guam | 13.372435 | 144.773357 | 4/27/2023 | high quality | Guam Department of Agriculture, Division of Aquatic and Wildlife Resources |
| YPM IZ 110605 | B1 | SW Pacific | Queensland | -28.089089 | 153.453921 | 2/3/2023 | high quality | Queensland Government General Fisheries Permit #209751; NSW Department of Primary Industries Scientific Collection Permit # P18/0012-1.0 |
| YPM IZ 110612 | B1 | SW Pacific | New South Wales | -30.745087 | 152.995494 | 3/26/2023 | high quality | Queensland Government General Fisheries Permit #209751; NSW Department of Primary Industries Scientific Collection Permit # P18/0012-1.0 |
| YPM IZ 110613 | B1 | SW Pacific | New South Wales | -30.745087 | 152.995494 | 3/26/2023 | high quality | Queensland Government General Fisheries Permit #209751; NSW Department of Primary Industries Scientific Collection Permit # P18/0012-1.0 |

Table 1: Specimen data. *(continued)*

| ID | cluster | ocean region | location | latitude | longitude | date collected | quality status | permit |
|---|---|---|---|---|---|---|---|---|
| YPM IZ 110616 | B1 | SW Pacific | New South Wales | -30.881192 | 153.035604 | 3/26/2023 | high quality | Queensland Government General Fisheries Permit #209751; NSW Department of Primary Industries Scientific Collection Permit # P18/0012-1.0 |
| YPM IZ 110631 | A | NW Atlantic | Northeast US | 41.346250 | -71.678310 | 7/14/2023 | high quality | RIDEM- Division of Marine Fisheries, 855, Type:1; RI: Scientific Collectors Permit #855 |
| YPM IZ 110633 | A | NW Atlantic | Northeast US | 41.377020 | -71.527820 | 7/15/2023 | high quality | RIDEM- Division of Marine Fisheries, 855, Type:1; RI: Scientific Collectors Permit #855 |
| YPM IZ 110667 | A | NW Atlantic | Northeast US | 40.857420 | -71.779920 | 7/20/2023 | high quality | NY: Scientific License #1427 |
| YPM IZ 110694 | A | NW Atlantic | Northeast US | 41.344990 | -70.836797 | 7/26/2023 | high quality | RIDEM- Division of Marine Fisheries, 855, Type:1; RI: Scientific Collectors Permit #855 |
| YPM IZ 110716 | A | NE Atlantic | Spain | 36.458090 | -6.252680 | 4/2/2013 | high quality | |
| YPM IZ 110726 | A | NE Atlantic | Spain | 36.458090 | -6.252680 | 4/2/2013 | high quality | |
| YPM IZ 110738 | A | NE Atlantic | Spain | 36.458090 | -6.252680 | 4/2/2013 | high quality | |
| YPM IZ 110777 | B1 | Central Pacific | Hawai'i | 21.400790 | -157.734500 | 7/19/2023 | high quality | |
| YPM IZ 110780 | B1 | Central Pacific | Hawai'i | 21.400790 | -157.734500 | 7/19/2023 | high quality | |
| YPM IZ 110785 | B1 | Central Pacific | Hawai'i | 21.400790 | -157.734500 | 7/21/2023 | high quality | |
| YPM IZ 110786 | B1 | Central Pacific | Hawai'i | 21.400790 | -157.734500 | 7/28/2023 | high quality | |
| YPM IZ 110828 | C1 | SE Pacific | Chile | -29.972700 | -71.360900 | 7/12/2023 | high quality | |
| YPM IZ 110829 | C1 | SE Pacific | Chile | -29.972700 | -71.360900 | 7/13/2023 | high quality | |
| YPM IZ 110830 | B1 | SW Atlantic | Uruguay | -33.751600 | -53.380100 | 2/28/2014 | high quality | |
| YPM IZ 110842 | A | NE Atlantic | Azores | 38.524769 | -28.625946 | 7/18/2019 | high quality | Secretaria Regional do Mar, Ciência e Tecnologia dos Açores, AMP/2018/021 |
| YPM IZ 110843 | A | NE Atlantic | Azores | 37.818604 | -25.543635 | 5/22/2020 | high quality | Secretaria Regional do Mar, Ciência e Tecnologia dos Açores, AMP/2018/021 |
| YPM IZ 110844 | A | NE Atlantic | Azores | 38.524769 | -28.625946 | 11/29/2019 | high quality | Secretaria Regional do Mar, Ciência e Tecnologia dos Açores, AMP/2018/021 |
| YPM IZ 110845 | A | NE Atlantic | Azores | 37.750488 | -25.624447 | 8/17/2020 | high quality | Secretaria Regional do Mar, Ciência e Tecnologia dos Açores, AMP/2018/021 |
| YPM IZ 110846 | A | NE Atlantic | Azores | 38.523756 | -28.623767 | 7/10/2020 | high quality | Secretaria Regional do Mar, Ciência e Tecnologia dos Açores, AMP/2018/021 |
| YPM IZ 110847 | A | NE Atlantic | Azores | 38.542865 | -28.618676 | 7/15/2020 | high quality | Secretaria Regional do Mar, Ciência e Tecnologia dos Açores, AMP/2018/021 |
| YPM IZ 110876 | A | Gulf of Mexico | Texas | 29.260000 | -94.840000 | 3/19/2017 | high quality | |
| YPM IZ 110877 | B2 | Gulf of California | Mexico | 27.245000 | -111.500000 | 6/6/2007 | high quality | |
| YPM IZ 110878 | B2 | Gulf of California | Mexico | 27.326667 | -111.480000 | 6/9/2006 | high quality | |
| YPM IZ 110880 | A | NW Atlantic | Northeast US | 41.149290 | -71.481140 | 8/23/2023 | high quality | RIDEM- Division of Marine Fisheries, scientific collectors permit:855 |
| YPM IZ 110881 | B1 | Central Pacific | Hawai'i | 19.666667 | -156.083333 | 4/10/2023 | high quality | |
| YPM IZ 110882 | B1 | Central Pacific | Hawai'i | 19.666667 | -156.083333 | 4/10/2023 | high quality | |
| YPM IZ 110883 | B1 | Central Pacific | Hawai'i | 19.666667 | -156.083333 | 4/10/2023 | high quality | |
| YPM IZ 110906 | B1 | Central Pacific | NW Hawaiian Islands | 26.723056 | -164.766667 | 8/23/2023 | high quality | Papahānaumokuākea Marine National Monument, PMNM-2023-001 |

Table 1: Specimen data. *(continued)*

| ID | cluster | ocean region | location | latitude | longitude | date collected | quality status | permit |
|---|---|---|---|---|---|---|---|---|
| YPM IZ 110907 | B1 | Central Pacific | NW Hawaiian Islands | 26.080000 | -176.470000 | 8/22/2023 | high quality | Papahānaumokuākea Marine National Monument, PMNM-2023-001 |
| YPM IZ 110925 | B1 | NW Pacific | Japan | 28.404780 | 129.454826 | 2/4/2023 | high quality | |
| YPM IZ 110926 | B1 | NW Pacific | Japan | 35.309561 | 139.478820 | 4/10/2023 | high quality | |
| YPM IZ 110930 | B1 | NW Pacific | Japan | 35.309561 | 139.478820 | 4/30/2023 | high quality | |
| YPM IZ 110931 | B2 | NW Pacific | Japan | 35.309561 | 139.478820 | 5/19/2023 | high quality | |
| YPM IZ 110933 | B1 | NW Pacific | Japan | 35.309701 | 139.478831 | 7/20/2023 | high quality | |
| YPM IZ 110937 | B1 | NW Pacific | Japan | 35.309561 | 139.478820 | 7/21/2023 | high quality | |
| YPM IZ 110938 | B1 | NW Pacific | Japan | 35.309701 | 139.478831 | 9/22/2023 | high quality | |
| YPM IZ 110943 | B2 | NW Pacific | Japan | 35.134935 | 140.284606 | 5/19/2023 | high quality | |
| YPM IZ 110972 | A | NE Atlantic | Ireland | 51.83707 | -10.19556 | 10/8/2023 | high quality | |
| YPM IZ 111006 | B1 | W Indian | South Africa | -27.539919 | 32.678650 | 8/8/2023 | high quality | Department of Fisheries, Forestry and Environment, amend RES2023-78 |
| YPM IZ 111007 | B1 | W Indian | South Africa | -27.539919 | 32.678650 | 8/8/2023 | high quality | Department of Fisheries, Forestry and Environment, amend RES2023-78 |
| YPM IZ 111008 | B1 | W Indian | South Africa | -27.539919 | 32.678650 | 8/8/2023 | high quality | Department of Fisheries, Forestry and Environment, amend RES2023-78 |
| YPM IZ 111009 | B1 | W Indian | South Africa | -27.539919 | 32.678650 | 8/8/2023 | high quality | Department of Fisheries, Forestry and Environment, amend RES2023-78 |
| YPM IZ 111010 | C1 | W Indian | South Africa | -27.539919 | 32.678650 | 8/8/2023 | high quality | Department of Fisheries, Forestry and Environment, amend RES2023-78 |
| YPM IZ 111012 | A | NW Atlantic | Bermuda | 32.307800 | -64.750500 | 3/8/2022 | high quality | Bermuda Special Permit no. SP200501 |
| YPM IZ 111013 | A | NW Atlantic | Bermuda | 32.307800 | -64.750500 | 3/8/2022 | high quality | Bermuda Special Permit no. SP200501 |
| YPM IZ 111014 | A | NW Atlantic | Bermuda | 32.307800 | -64.750500 | 3/9/2022 | high quality | Bermuda Special Permit no. SP200501 |
| YPM IZ 111016 | A | NW Atlantic | Bermuda | 32.307800 | -64.750500 | 3/9/2022 | high quality | Bermuda Special Permit no. SP200501 |
| YPM IZ 111017 | A | NW Atlantic | Bermuda | 32.307800 | -64.750500 | 3/9/2022 | high quality | Bermuda Special Permit no. SP200501 |
| YPM IZ 111018 | A | NW Atlantic | Bermuda | 32.307800 | -64.750500 | 3/9/2022 | high quality | Bermuda Special Permit no. SP200501 |
| YPM IZ 111156 | C1 | SW Pacific | Tasman Sea | -35.747000 | 154.330000 | 10/16/2023 | high quality | |
| YPM IZ 111157 | C1 | SW Pacific | Tasman Sea | -35.747000 | 154.330000 | 10/16/2023 | high quality | |
| YPM IZ 111158 | C1 | SW Pacific | Tasman Sea | -35.747000 | 154.330000 | 10/16/2023 | high quality | |
| YPM IZ 111159 | C1 | SW Pacific | Tasman Sea | -35.747000 | 154.330000 | 10/16/2023 | high quality | |
| YPM IZ 111160 | C1 | SW Pacific | Tasman Sea | -35.747000 | 154.330000 | 10/16/2023 | high quality | |
| YPM IZ 111163 | B1 | SW Pacific | New South Wales | -33.947500 | 151.257361 | 3/18/2024 | high quality | |
| YPM IZ 111207 | C2 | SW Pacific | New Zealand | -36.895150 | 174.443889 | 3/15/2024 | high quality | |
| YPM IZ 111208 | C2 | SW Pacific | New Zealand | -36.895150 | 174.443889 | 3/15/2024 | high quality | |
| YPM IZ 111209 | C2 | SW Pacific | New Zealand | -36.895150 | 174.443889 | 3/15/2024 | high quality | |
| YPM IZ 111210 | C2 | SW Pacific | New Zealand | -36.895150 | 174.443889 | 3/15/2024 | high quality | |
| NIWA 173304 | C2 | SW Pacific | New Zealand | -36.895150 | 174.443889 | 3/15/2024 | high quality | |
| NIWA 173305 | C2 | SW Pacific | New Zealand | -36.895150 | 174.443889 | 3/15/2024 | high quality | |

#### Specimen photos

The following arrays of photos show specimens upon collection (left) and after ethanol fixation (right), when both views are available.

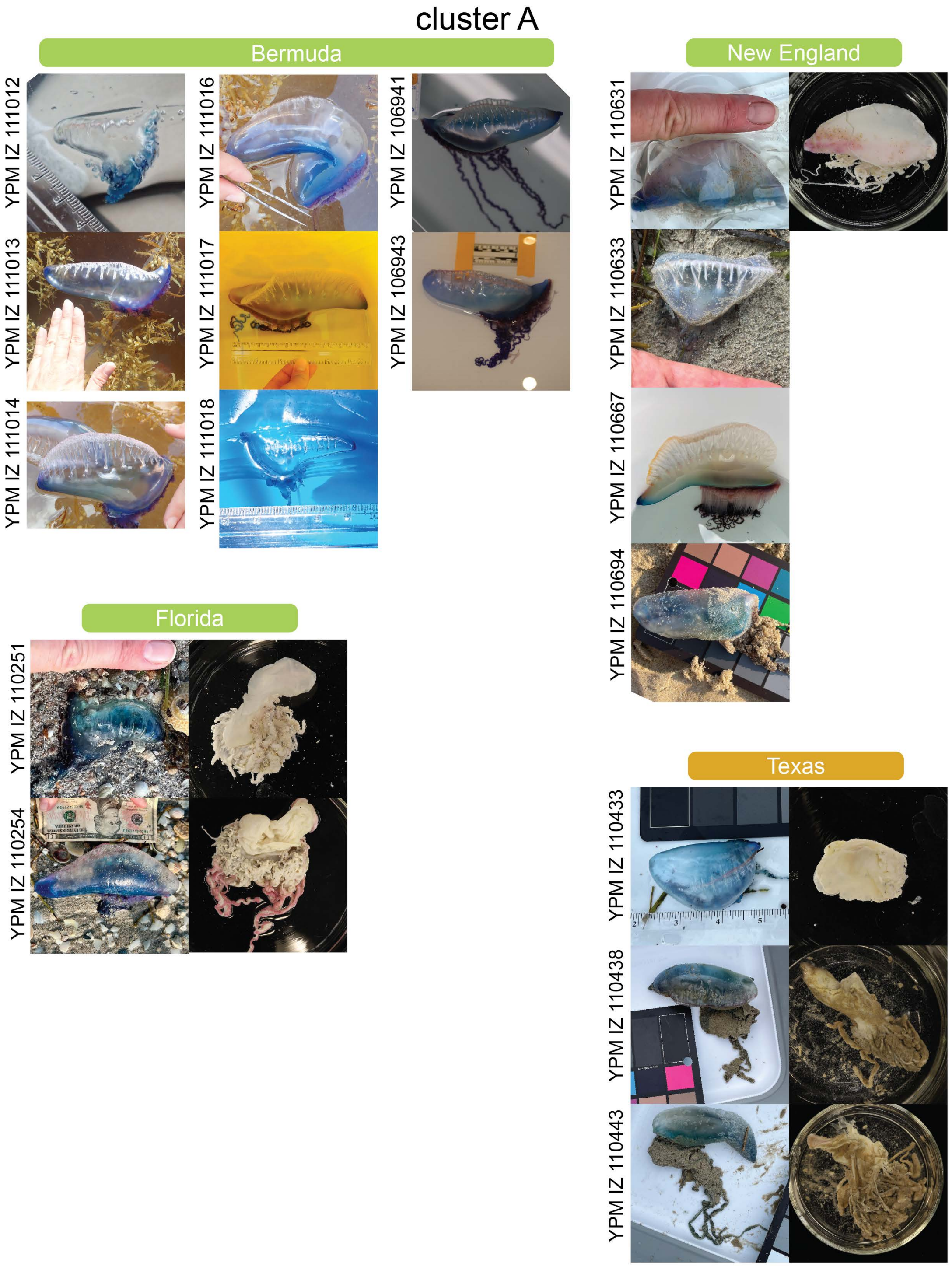


Figure 1: Specimen photos: cluster A from the NW Atlantic.

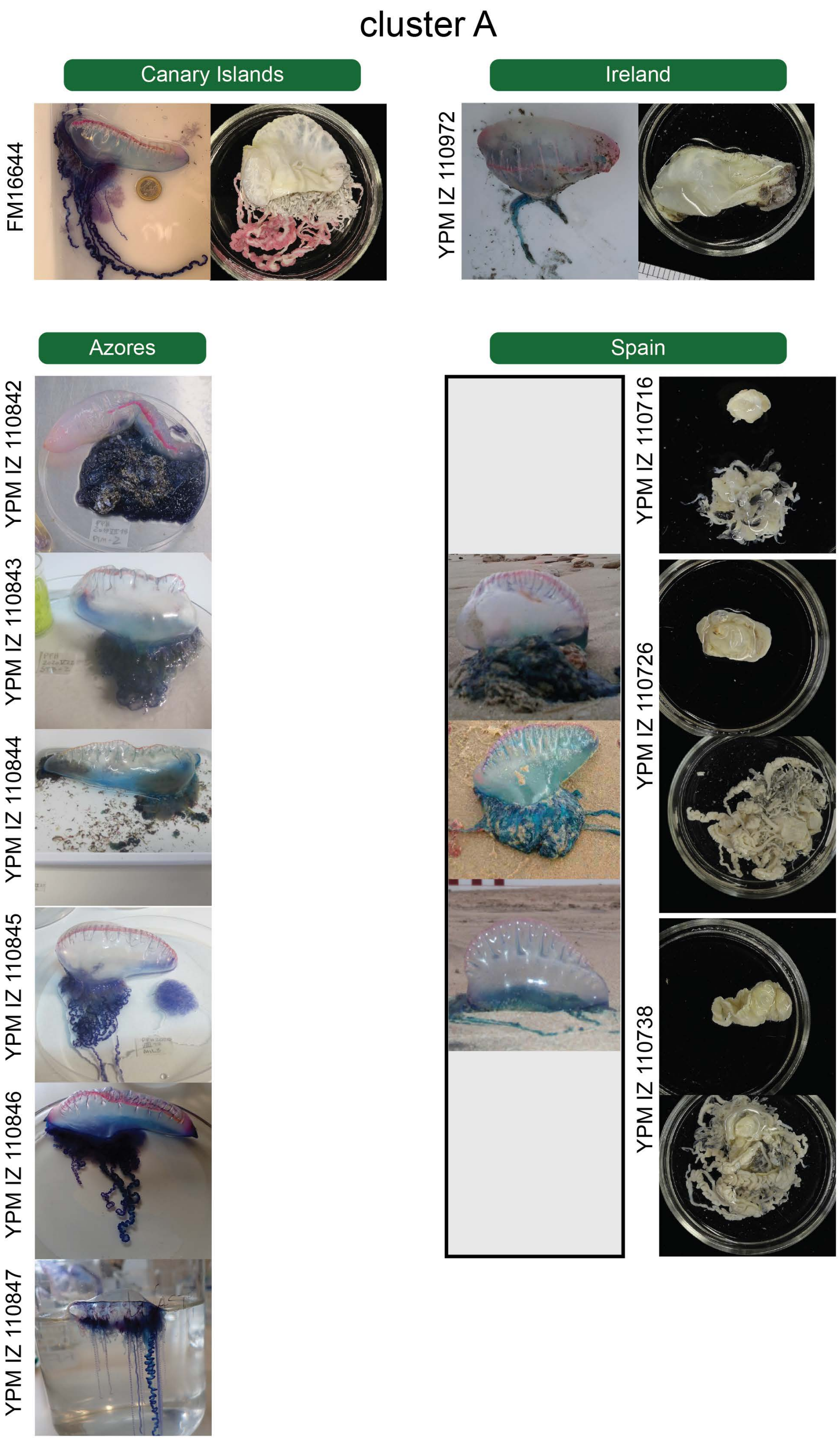


Figure 2: Specimen photos: cluster A from the NE Atlantic.

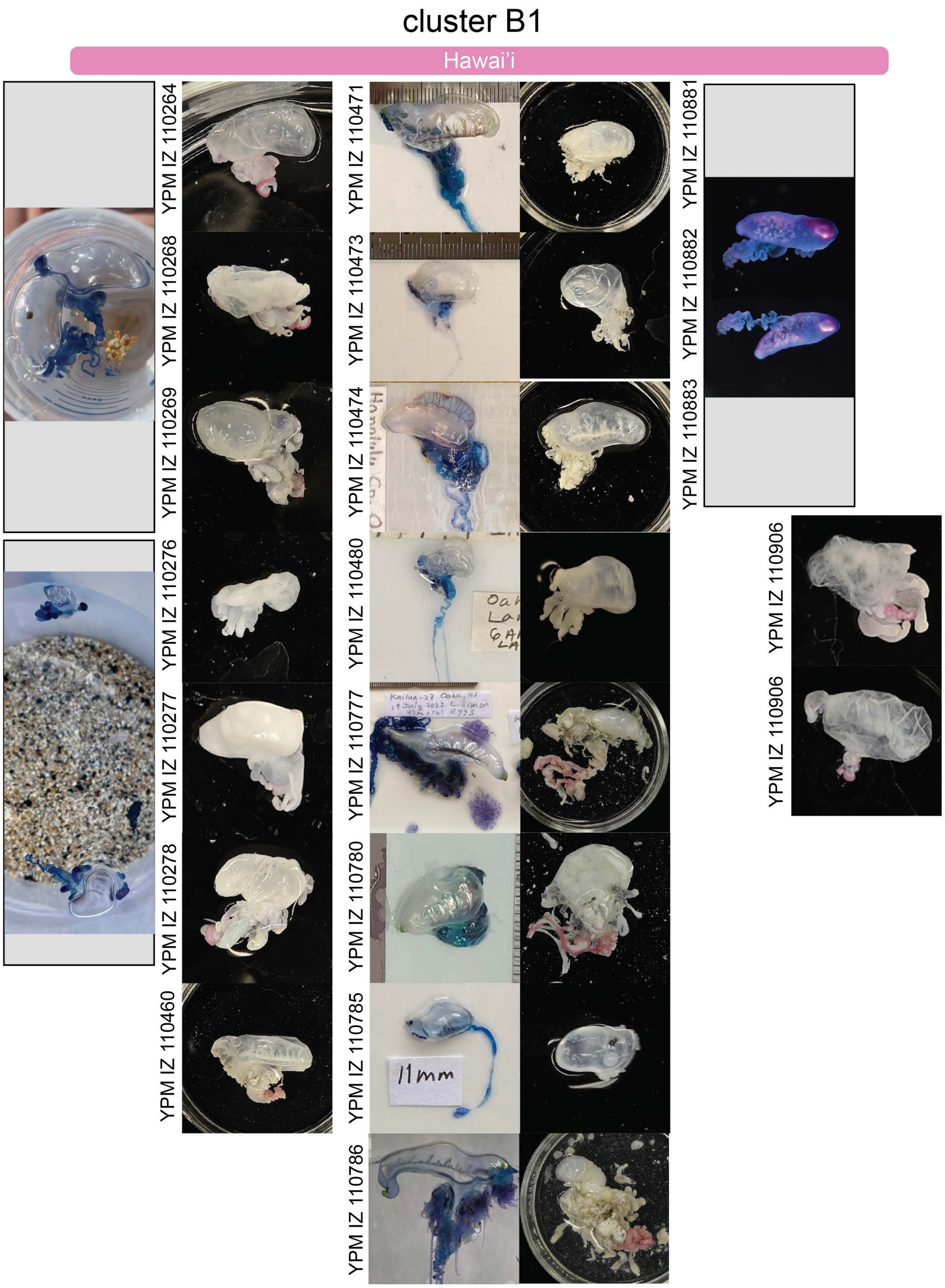


Figure 3: Specimen photos: cluster B1 from Hawai'i and the NW Hawaiian Islands.

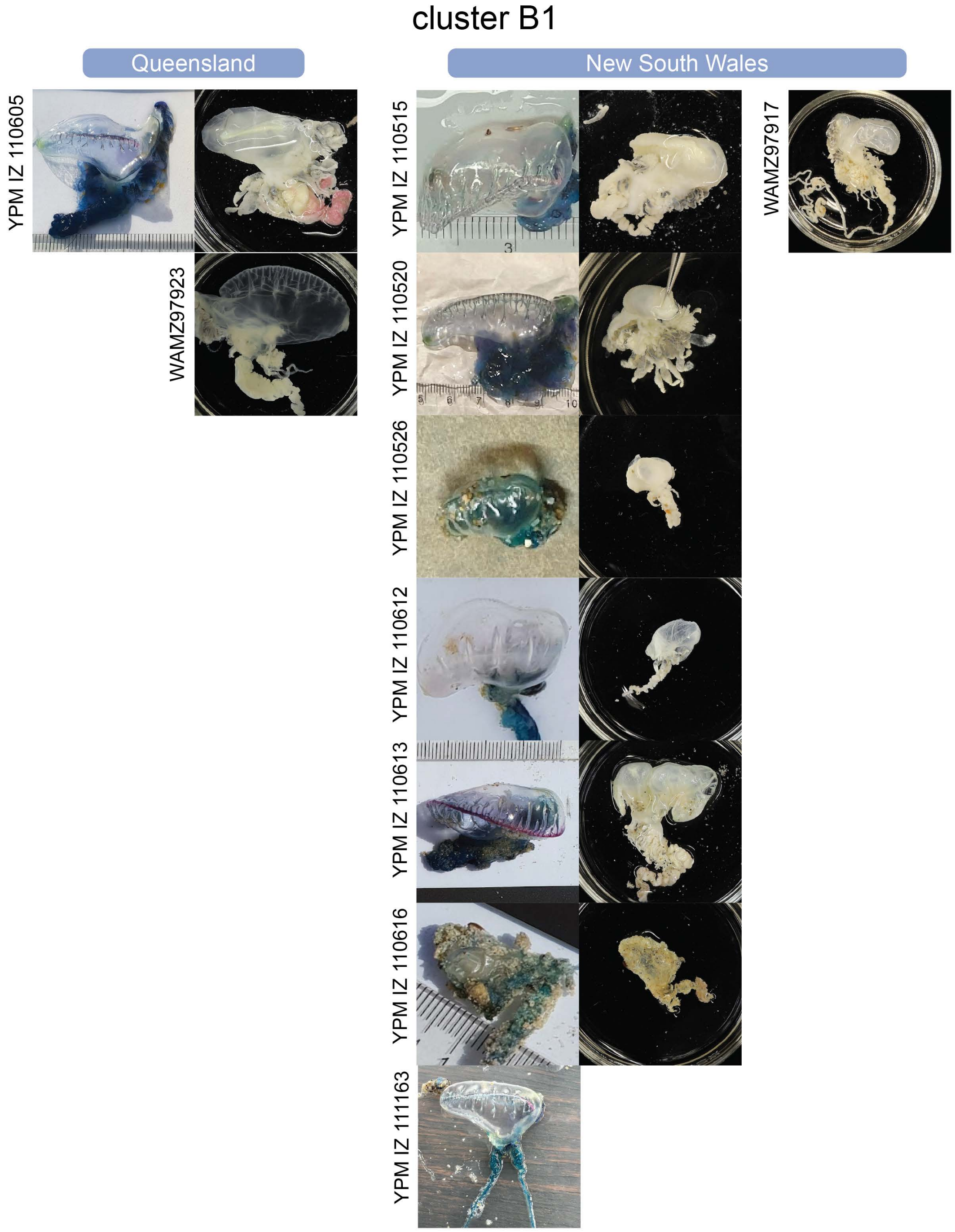


Figure 4: Specimen photos: cluster B1 from E Australia.

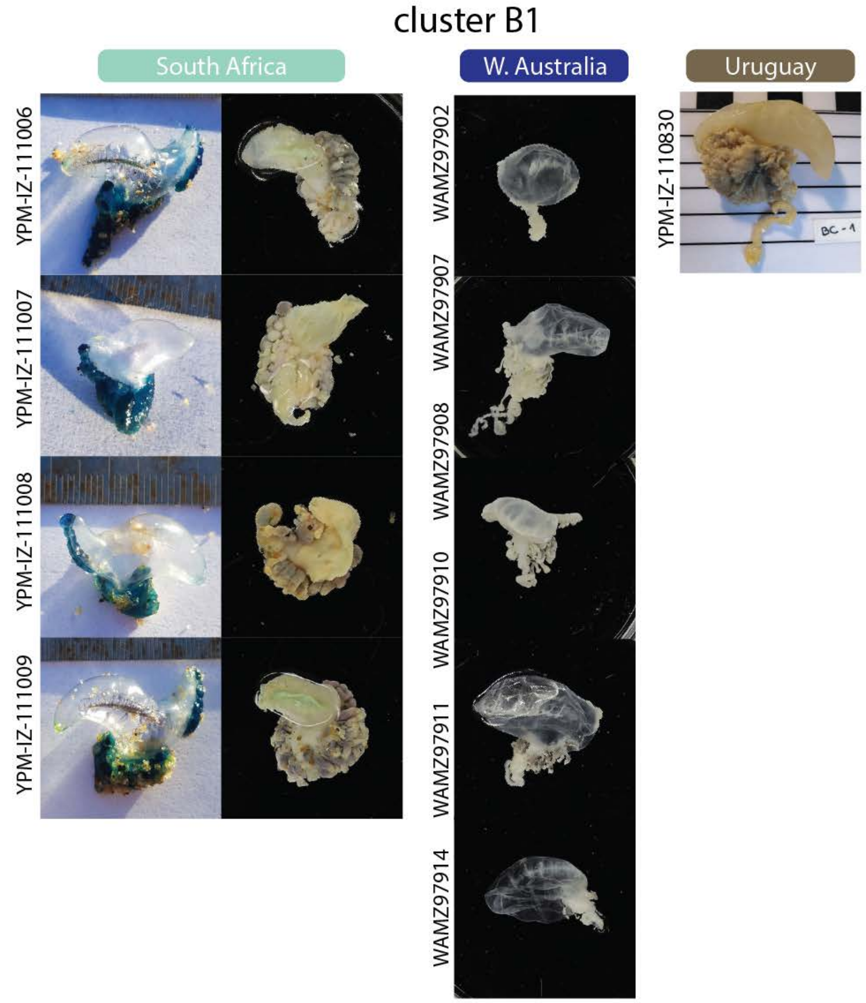


Figure 5: Specimen photos: cluster B1 from the S Indian and S Atlantic.

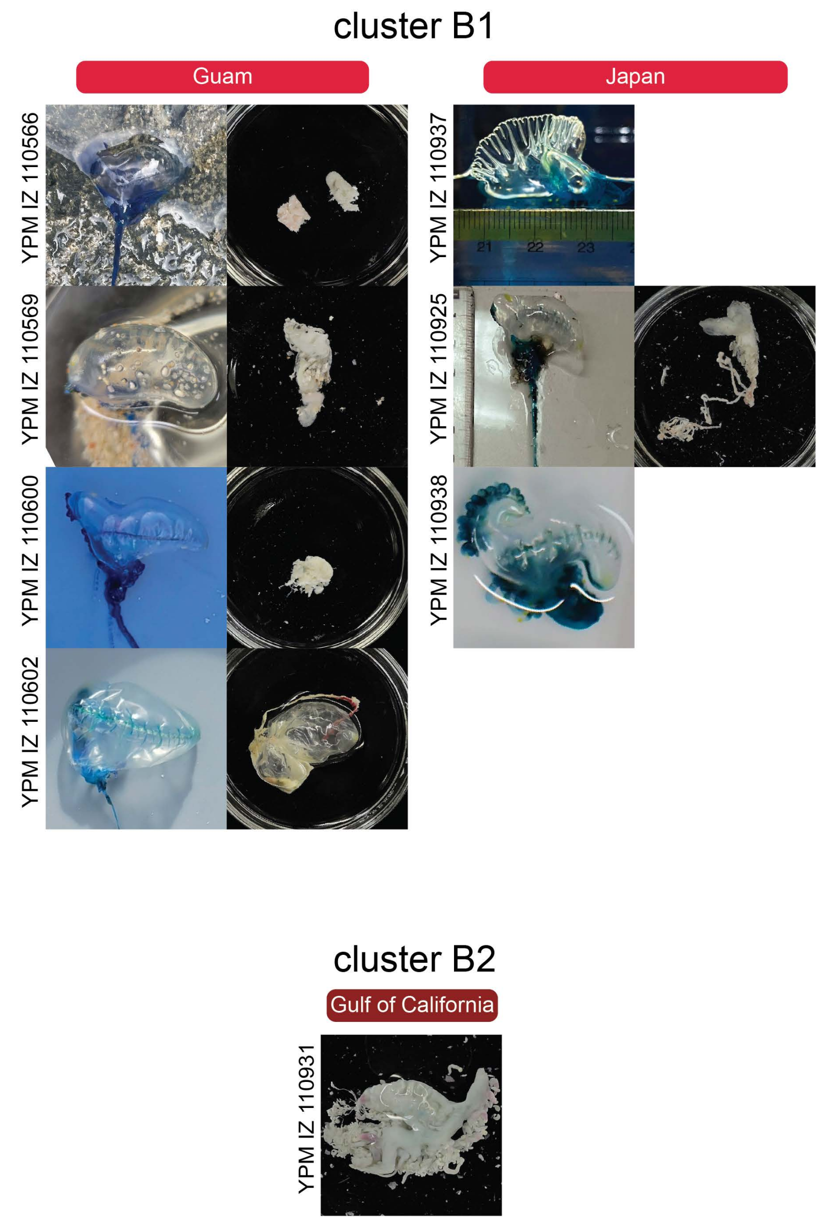


Figure 6: Specimen photos: cluster B1 from the NW Pacific, and cluster B2.

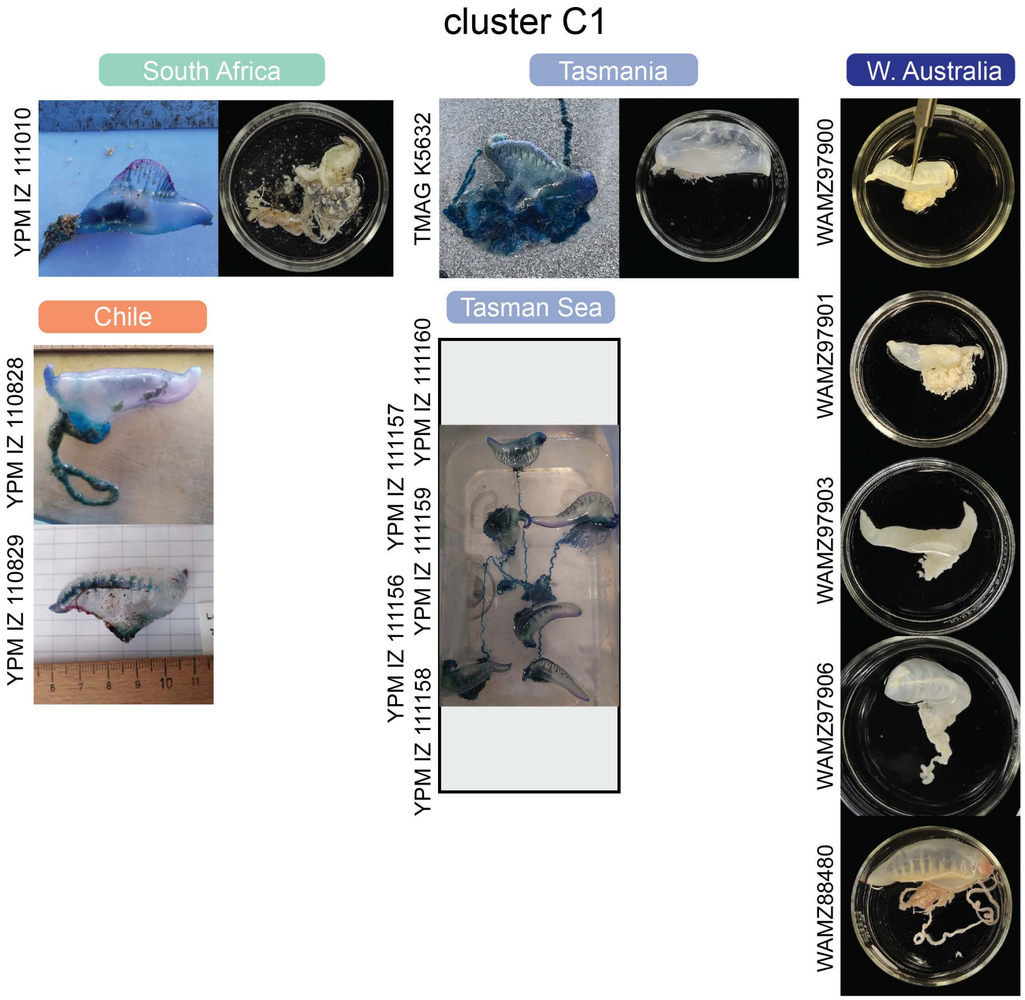


Figure 7: Specimen photos: cluster C1.

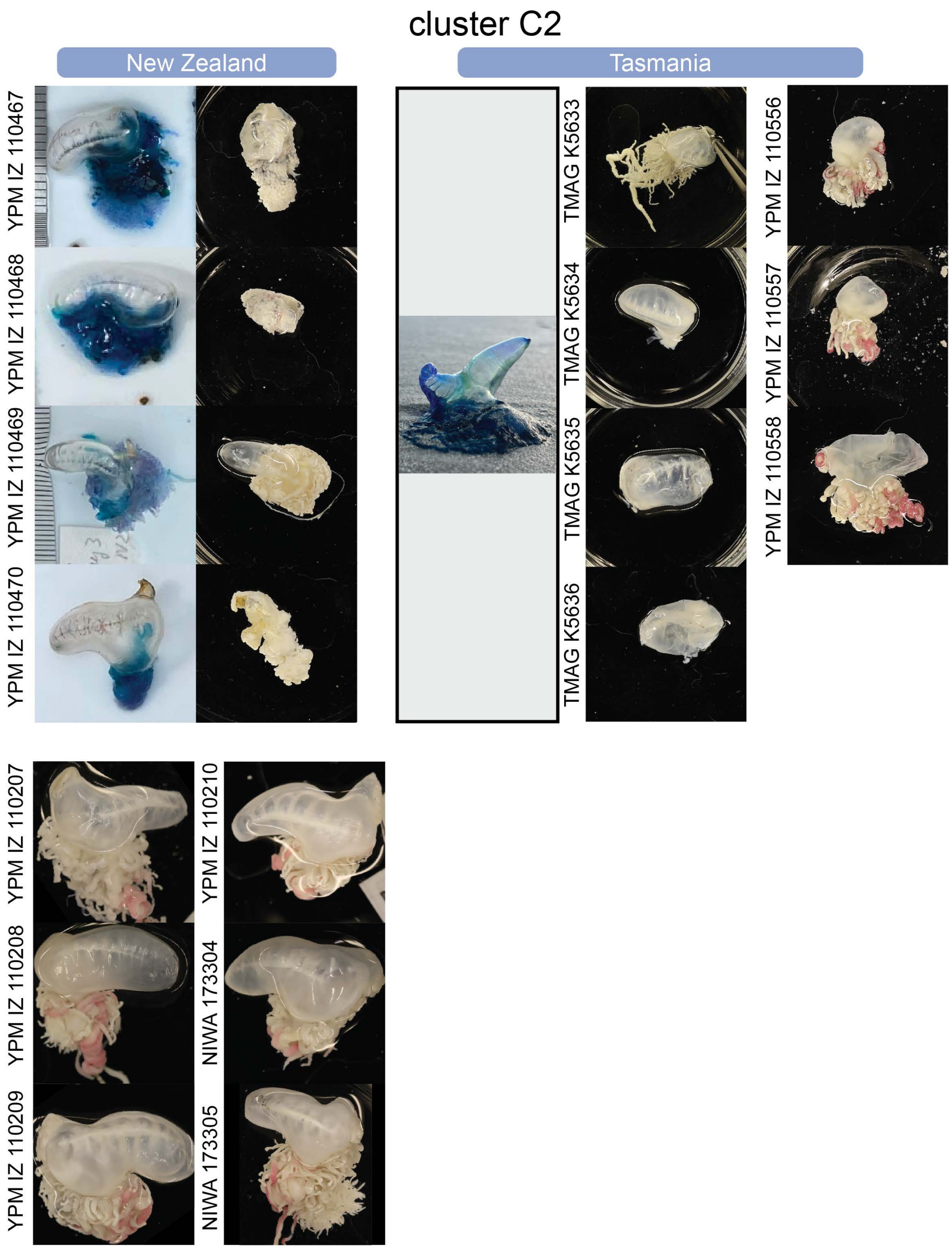


Figure 8: Specimen photos: cluster C2.
